# Transient Maternal IL-6 boosts glutamatergic synapses and disrupts hippocampal connectivity in the offspring

**DOI:** 10.1101/2020.11.02.364356

**Authors:** Filippo Mirabella, Genni Desiato, Sara Mancinelli, Giuliana Fossati, Marco Rasile, Raffaella Morini, Marija Markicevic, Christina Grimm, Clara Amegandjin, Alberto Termanini, Clelia Peano, Paolo Kunderfranco, Graziella di Cristo, Valerio Zerbi, Simona Lodato, Elisabetta Menna, Michela Matteoli, Davide Pozzi

**Affiliations:** Humanitas University, Department of Biomedical Science, Pieve Emanuele, Milan, 20090, Italy; Humanitas Clinical and Research Center - IRCCS, Rozzano, Milan, 20089, Italy; Neuroscience Center Zurich, ETH Zurich and University of Zurich, Zurich, 8057, Switzerland; Department of Neurosciences, Université de Montréal, Montréal, Québec, Canada; CHU Sainte-Justine Research Center, Montréal, Québec, Canada; Bioinformatic Unit, Humanitas Clinical and Research Center, Rozzano, Milan, 20089, Italy; Institute of Genetic and Biomedical Research, UoS Milan, National Research Council, Rozzano, Milan, 20089, Italy; Genomic Unit, Humanitas Clinical and Research Center, Rozzano, Milan, 20089, Italy; Institute of Neuroscience - National Research Council, Milan, 20139, Italy; Neural Control of Movement Lab, Department of Health Sciences and Technology, ETH Zürich, Zurich, 8057 Switzerland

**Keywords:** Neurodevelopmental Disorder, Synaptic development, IL-6, STAT3, RGS4, Pro-inflammatory cytokines, Neuroinflammation, Glutamatergic Transmission, Maternal Immune Activation, Brain Connectivity

## Abstract

Early prenatal inflammatory conditions are thought to represent a risk factor for different neurodevelopmental disorders, with long-term consequences on adult brain connectivity. Here we show that a transient IL-6 elevation, occurring at vulnerable stages of early neurodevelopment, directly impacts brain developmental trajectories through the aberrant enhancement of glutamatergic synapses and overall brain hyper-connectivity. The IL6-mediated boost of excitatory synapse density results from the neuron-autonomous, genomic effect of the transcription factor STAT3 and causally involves the activation of RGS4 gene as a candidate downstream target. The STAT3/RGS4 pathway is also activated in neonatal brains as a consequence of maternal immune activation protocols mimicking a viral infection during pregnancy. By demonstrating that prenatal IL-6 elevations result in aberrant synaptic and brain connectivity through the molecular players identified, we provide a mechanistic framework for the association between prenatal inflammatory events and brain neurodevelopmental disorders.

## Introduction

The formation of synapses during the development of the central nervous system (CNS) represents a critical process which ensures a proper brain connectivity patterns at adult stages (Lu et al, 2009; Williams et al, 2010). The centrality of this process has been highlighted by the evidence that several neurodevelopmental disorders, including Autism Spectrum Disorders (ASD) (Bourgeron, 2009; Delorme et al, 2013) and Schizophrenia (Hall et al, 2015; Owen et al, 2016), are characterized by defects in the formation, maturation and maintenance of synaptic contacts (Melom & Littleton, 2011; Penzes et al, 2011), resulting in altered brain development (Courchesne et al, 2007; Supekar et al, 2013)

The dynamic of synapse formation is a complex and hierarchically regulated event in which both intrinsic and extrinsic factors act together to ensure a proper brain connectivity (McAllister, 2007) and a correct excitatory/inhibitory (E/I) balance (Cline, 2005; Gatto & Broadie, 2010). Beyond the highly specialized genetic program, which allows the precise temporal expression of neuronal genes involved in synaptogenesis (Shen & Scheiffele, 2010), recent evidences highlighted the critical contribution of environmental factors in the modulation of synapse formation (Grabrucker, 2012). Among them, inflammatory states occurring at early stages of neuronal development have been recognized as the main environmental insult which may negatively affect the entire brain developmental trajectory. Indeed, epidemiological and experimental studies indicate a clear association between inflammation during pregnancy and risk of neurodevelopmental disorders in the progeny (Bergink et al, 2014; Knuesel et al, 2014; Li et al, 2009; Onore et al, 2012; Potvin et al, 2008). Although the underlying mechanisms are still unknown, soluble immune molecules released upon inflammatory states are thought to be key players in this process (Bauer et al, 2007; Deverman & Patterson, 2009; McAfoose & Baune, 2009).

A molecule widely engaged during inflammation is interleukine-6 (IL-6). IL-6 is a pleiotropic proinflammatory cytokine that exerts several actions on the mature nervous system (Balschun et al, 2004; Erta et al, 2012), modulating a plethora of brain processes including energy homeostasis (Timper et al, 2017; Wallenius et al, 2002), adult neurogenesis (Monje et al, 2003; Vallieres et al, 2002), and axonal regeneration upon neuronal damage (Cafferty et al, 2004; Leibinger et al, 2013a; Leibinger et al, 2013b; Pieraut et al, 2011). Besides these effects, the cytokine also plays key actions during brain development. Indeed, mice embryos prenatally exposed to IL-6 display behavioral defects at adult stages (Choi et al, 2016; Shin Yim et al, 2017; Smith et al, 2007b), while a tight association between elevated IL-6 levels in the pregnant mother and altered brain connectivity and working memory in the newborns has been reported in humans (Rudolph et al, 2018; Spann et al, 2018). These evidences strongly suggest that an early IL-6 elevation can deeply affect neuronal development, leading to behavioral abnormalities. However, whether this phenomenon might be linked to a long-lasting alteration of synaptic formation is still undefined.

Here we describe an hippocampal pro-synaptogenic effect of IL-6 which specifically involves glutamatergic synapses. The increased formation of excitatory synapses persists at mature stages and is associated with brain hyper-connectivity. We also demonstrate that this process depends on the activation of Signal transducer and activator of transcription-3 (STAT3) and involves its downstream target gene Regulator of G protein Signaling 4 (RGS4). Therefore, a transient increase of IL-6, as a consequence of inflammatory processes occurring at early phases of neuronal development, is sufficient to disarrange the entire process of excitatory synaptogenesis resulting in an abnormal brain connectivity in the adulthood.

## Results

### Transient prenatal IL-6 elevation enhances hippocampal glutamatergic synapses in the adulthood

We first aimed to assess whether transient prenatal IL-6 elevations impact the density of synaptic contacts in the offspring at postnatal stages. To this aim, a single intraperitoneal (i.p) injection of either 5 μg of IL-6 (Choi et al, 2016; Gallagher et al, 2013; Smith et al, 2007b) or vehicle (as control), was administered to pregnant mice at gestational day 15 (GD15) (Figure S1A), when hippocampus is already formed (Urban & Guillemot, 2014), the neurogenesis is peaking whereas synaptogenesis and astrogenesis have not started yet (Angevine, 1965; Finlay & Darlington, 1995; Reemst et al, 2016).

The number of excitatory and inhibitory synaptic puncta was evaluated in the CA1 radiatum area of mice at postnatal day (P) 15 through immunofluorescence analysis of both glutamate (V-glut) and GABA (V-gat) vesicular transporters (Figure 1A). We found that the area of V-glut, but not V-gat, positive puncta was significantly enhanced in the hippocampus of mice exposed to IL-6 at prenatal stages (Figure 1B). To evaluate whether the selective increase of glutamatergic synaptic inputs was accompanied by functional changes, glutamatergic and GABAergic synaptic basal transmission were simultaneously evaluated at single cell level by whole cell patch-clamp recording of pyramidal neurons in the CA1 hippocampal region (Figure 1C). We found a substantial increase in the frequency of miniature excitatory post synaptic currents (mEPSCs) in the hippocampi of mice prenatally exposed to IL-6 as compared to vehicle (Figure 1D), whereas miniature excitatory post synaptic currents (mIPSCs) were not altered (Figure 1E), resulting in an excitatory/inhibitory (E/I) imbalance of neurotransmission (Figure 1F)

**Figure 1.**
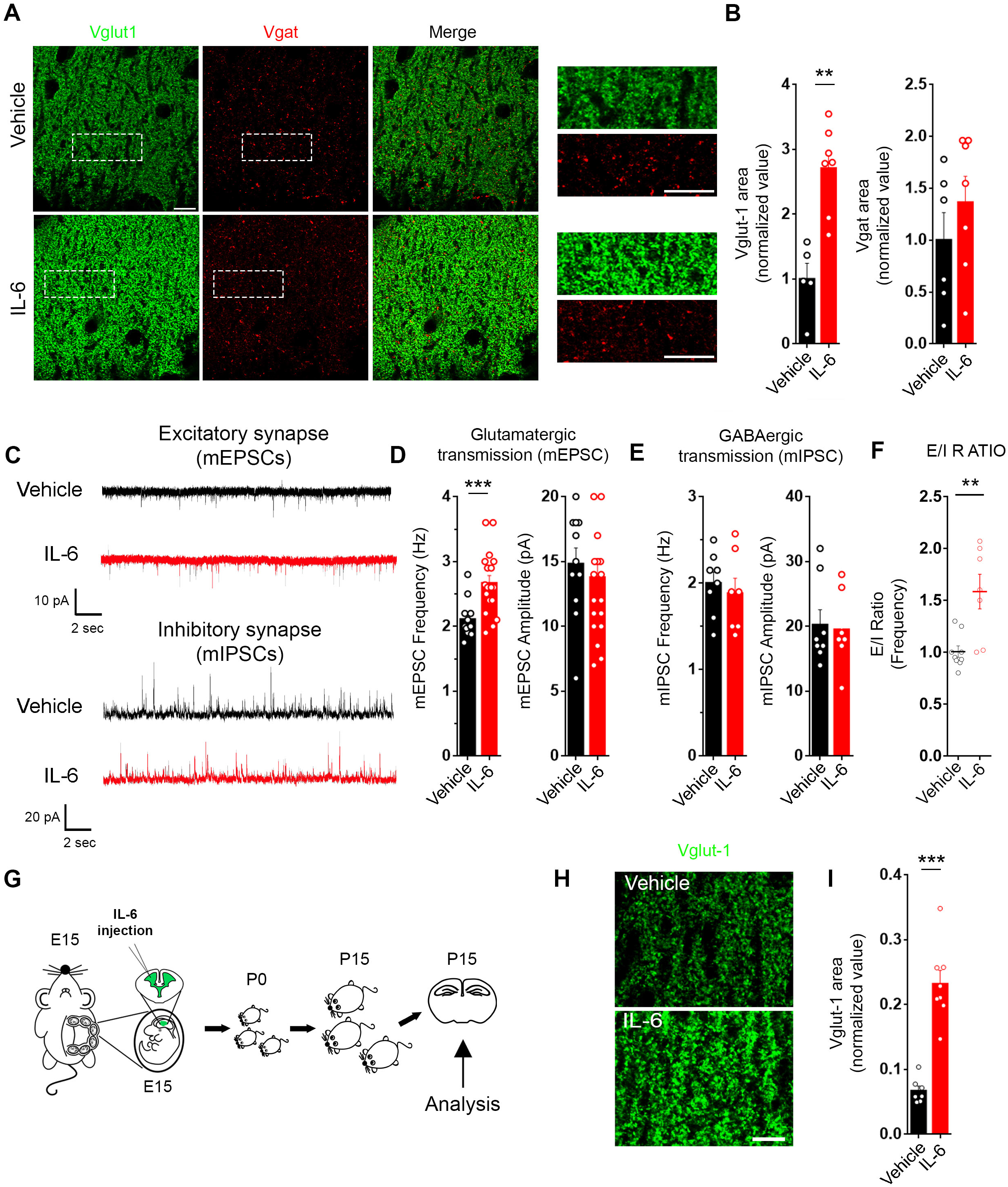
Transient prenatal exposure of embryos to IL-6 at E15 increases Glutamatergic synapses in the offspring at post-natal stages. **(A)** Immunofluorescence analysis of Glutamatergic and GABAergic synapses identified as Vglut-1 (in green) or V-gat (in red) positive puncta respectively performed in the stratum radiatum of CA1 hippocampal region of offspring derived from Vehicle-(Vehicle) or IL-6 treated mother (IL-6). Scale bar 5 μm. Right panels, high magnification image relative to the dotted square. **(B)** Quantitative analysis of Vglut-1 (vehicle n= 5 mice; IL-6 n=7 mice. 3 independent experiments) and V-gat (vehicle n= 6 mice; IL-6 n=7 mice. 3 independent experiments) area in the two conditions. Mann-Whitney test. **p= 0,0025. **(C)** Electrophysiological traces of mEPSCs and mIPSCs recorded in CA1 pyramidal hippocampal neurons in acute brain slices established from P15 offspring embryonically exposed to Vehicle or IL-6. **(D-E)** Quantitative analysis of mEPSCs (vehicle n= 12 cells, 4 mice; IL-6 n=18 cells, 5 mice. 3 independent experiments) and mIPSCs (vehicle n= 7 cells, 3 mice; IL-6 n=8 cells, 3 mice. 3 independent experiments) frequency and amplitude. Mann-Whitney test ***p=0,0009. **(F)** quantitative analysis of E/I ratio calculated as ratio between mEPSCs and mIPSCs frequency recorded at single cell level. (vehicle n= 9 cells, 3 mice; IL-6 n=7 cells, 3 mice. 3 independent experiments) Mann-Whitney test **p=0,0032 **(G)** Schematic representation of the experimental procedure: intraventricular injection of vehicle or IL-6 (10ng/embryo) were performed in E15 embryos and the offspring were analyzed at post-natal day 15 by evaluating excitatory synapses. **(H)** Immunofluorescence analysis of Vglut-1 and V-gat positive puncta respectively performed in the stratum radiatum of CA1 hippocampal region of P15 offspring exposed to vehicle-(vehicle) or IL-6 via intraventricular injection. Scale bar 10 μm. **(I)** Quantitative analysis of Vglut-1 area in CA1 hippocampal region in the two conditions. (vehicle n= 7 injected mice; IL-6 n= 8 injected mice). Mann-Whitney test. ***p=0,0003.

Since maternal IL-6 elevation during pregnancy triggers the activation of peripheral cells, immune molecules and gut microbiota in the mother (Choi et al, 2016; Kim et al, 2017; Shin Yim et al, 2017) which could indirectly be responsible for the increase of glutamatergic synapses, IL-6 or vehicle were injected intraventricularly (ICV) in the embryos at E15 (Figure 1G) to bypass any indirect signaling of the mother. Synapse quantitation in P15 offspring hippocampi (Figure 1H) revealed that, similarly to results obtained upon intraperitoneal injection, ICV administration of IL-6 promotes the increase of glutamatergic synapses (Figure. 1I).

### Prenatal IL-6 elevation disrupts hippocampal functional connectivity in the adulthood

The selective increase in excitatory, but not inhibitory, synapses in mice exposed to maternal IL-6 elevation was found to persist up to P30 (Figure S1B-C). An abnormal number of excitatory synapses and/or an altered E/I balance within local microcircuits, found in many models of neurodevelopmental disorders (Durand et al, 2007; Lee et al, 2015; Sala et al, 2001), is often associated with macroscale alterations in functional connectivity that can be detected with resting-state fMRI (Ajram et al, 2017; Filipello et al, 2018; Pagani et al, 2019; Zhou et al, 2019). Hence, we determined whether the transient maternal IL-6 elevation had an effect on brain connectivity in adulthood. We therefore acquired resting state-fMRI (rs-fMRI) scans in 16 mice (9 treated and 7 controls) at 14 weeks of age using a standardized pipelines for anesthesia control, data acquisition and preprocessing (Zerbi et al, 2015; Zerbi et al, 2018) (Figure 2A). To probe the existence of aberrant functional connections, the blood-oxygen-level-dependent (BOLD) time series were extracted from 165 regions of interest (ROIs) using the Allen’s Common Coordinate Framework and their connectivity couplings were measured using regularized Pearson’s correlation coefficients. Randomized permutation testing (5,000 permutations) revealed an overall hyper-connectivity phenotype of IL-6 mice compared to controls (Figure 2B). The spatial distribution of the hyper-connected edges was widespread (242 out of 2724 edges were identified as significantly hyper-connected at p<0.05) and the strongest contribution was given by hippocampal-midbrain, hippocampal-parietal, hippocampal-cortical subplate, prefrontal-parietal and somatomotor-thalamic connections among all (Figure 2C). Only 36 edges (1.3%) were found to be hypo-connected in the IL-6 group as compared to vehicle controls (Figure S1D).

**Figure 2:**
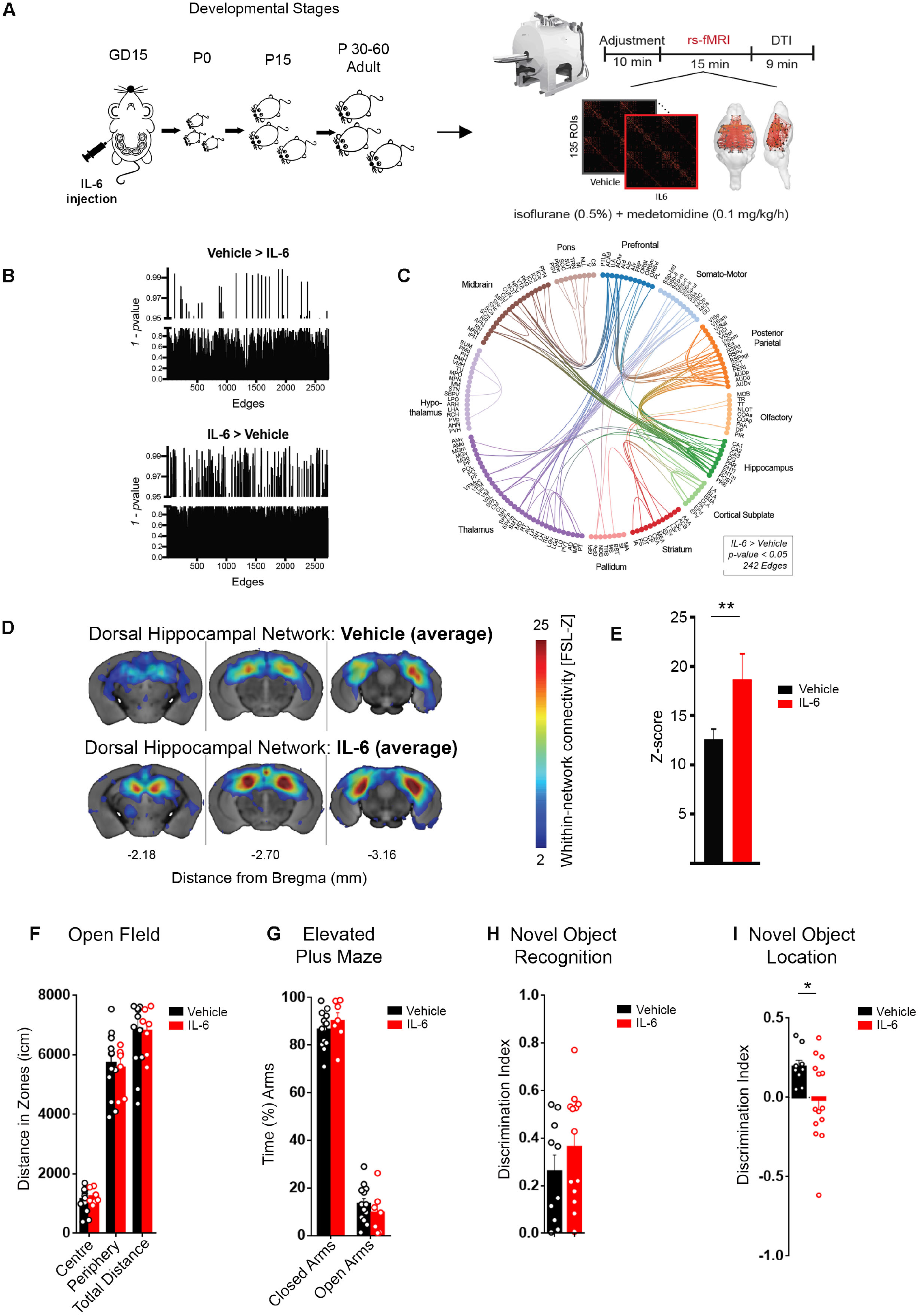
Offspring prenatally exposed to IL-6 show hyppocampal hyperconnettivity and altered spatial memory. **(A)** Schematic of the experimental set-up for MRI recordings. Adult offspring from mothers exposed to IL-6 or Vehicle at GD15 underwent a single MRI session to record resting-state fMRI and diffusion tensor imaging, under light anesthesia (isoflurane 0.5% + medetomidine 0.1 mg/kg/h). **(B)** Randomized non-parametric statistics of whole-brain functional *connectome* mapping indicates a shift toward hyper-connectivity in IL-6 mice **(C)** Circos-plot showing the anatomical location of hyper-connected edges (n=242) in IL-6 mice compared to vehicle-treated mice (*p*<0.05, uncorrected). (Vehicle n= 9 mice, IL-7 n= 7 mice. Three independent experiments) **(D)** Dual-regression analysis in 15 resting-state networks (RSNs) revels a significant increase in the Dorsal Hippocampal network strength in the IL-6 treated group compared to vehicle control mice (p=0.017, Bonferroni corrected). (**E)** Multivariate ANOVA, Bonferroni corrected across 15 RSNs. Bar plots represent mean ± SEM. **=*p*<0.01. (**F-I)** Behavioral evaluation of motor and cognitive functions. **(F-G)** Open field (F): Quantification of distance travelled separately in the center and periphery, and total distance (in cm) in an open field arena. Elevated plus maze **(G)**: quantification of the percentage of time spent in the closed and open arms of plus maze arena. Two-way Anova with Sidak’s multiple comparison *Post hoc* test. Vehicle n=13 male mice, IL-6 n=7 male mice, from 3 independent mothers. **(H-I) Discrimination** *index* values in novel object recognition (H) and in novel object location (I). Vehicle n=10 male mice, IL-6 n=13 male mice from 2 independent mothers. *p=0,0167 Unpaired t test with Welch’s correction.

Next, we examined whether the excessive excitatory neurotransmission by IL-6 exposure prompts large-scale resting-state network (RSN) reconfiguration. The connectivity strength within fifteen maximally-independent RSNs was measured using a dual regression approach as described elsewhere (Filippini et al, 2009) (for a complete list and spatial distribution of the networks please refer to our previous study (Zerbi et al, 2015)). Statistical analysis was conducted by comparing the connectivity-strength within all the voxels that constitute each RSNs. In the dorsal hippocampal network, the connectivity was significantly higher in IL-6 group compared to controls (Figure 2D,E). The temporal association network also showed moderate increases in connectivity in the IL-6 group, without reaching statistical significance (p=0.09). Conversely, a reduction in connectivity approaching significance was seen in the primary and secondary somatosensory network (p=0.066 and p=0.130, respectively (Figure S1E). None of the other networks were affected. These data suggest that excitatory neurotransmission after IL-6 exposure can be detected with rs-fMRI in the form of an increased synchronicity, especially within elements of the hippocampal network. The structural integrity of major axonal bundles was quantified by extracting fractional anisotropy (FA) values from seven whitematter structures as described (Zerbi et al, 2013b; Zerbi et al, 2019). All white matter tracts did not exhibit significant differences between IL-6 mice and vehicle controls (Figure S1F), indicating that prenatal IL-6 does not compromise the macroscopic characteristics of anatomical connections but rather impairs their function. In support of this, no major anatomical alterations were detected by Nissl staining in the cortex or hippocampus of mice exposed to IL-6 at E15 (Figure 2SA). Furthermore, analysis of cortical lamination in brains at P30 using antibodies for layer-specific markers including Special AT-rich sequence binding protein 2 (SATB2), which specifically labels commissural excitatory neurons in all cortical layers (Alcamo et al, 2008) and NeuroD2, pan-glutamatergic cortical neurons, did not reveal any defect in cortical architecture nor lamination in any experimental conditions (Figure 2SB,C). These data indicate that IL-6 exposure does not significantly alter the overall architecture of the cerebral cortex in the offspring. In addition, the brain of mature offspring (P30) did not display signs of astrogliosis and inflammation, as indicated by the lack of difference in the number and expression of the Glial Fibrillary Acidic Protein (GFAP) protein, a marker of astrocytes, and by the absence of altered number and morphology of Iba1-positive cells, a marker of microglia (Figure 2SC,D). Thus prenatal exposure of IL-6 does not lead to a chronic inflammatory state, nor astrogliosis, in the adult offspring.

Given the central role played by the hippocampus in specific forms of learning and memory, a derangement of hippocampal synaptic functioning and connectivity might be associated with an impairment of hippocampal-dependent behavioral task. A panel of tests was then employed to evaluate the possible occurrence of behavioral abnormalities (Figure 2F-I). We found that the prenatal exposure to IL-6 does not impinge the mice performance in the open field, elevated plus maze and novel object recognition tests (Figure 2F-G), indicating that locomotor activity, anxiety-like behavior and episodic memory were not compromised. However, when assayed through novel object location test, mice prenatally exposed to IL-6 showed a significantly lower discrimination index (Figure 2I), indicating a specific impairment of spatial memory.

These data demonstrate that a transient prenatal exposure to IL-6, in our experimental paradigm, results in increased glutamatergic synaptic contacts and enhanced excitatory neurotransmission, which associate with altered hippocampal-related functional connectivity and behavior, in the absence of major morphological defects and inflammatory signs.

### The pro-inflammatory cytokine IL-6 selectively enhances glutamatergic synaptogenesis through a direct action on neurons

We next aimed to gain more insights into the cellular and molecular mechanisms responsible for the cytokine effect. To investigate whether IL-6 enhances the intrinsic capacity of neurons to promote glutamatergic synapse formation, E18 embryos exposed to either IL-6 or vehicle at E15 were collected and employed to establish primary cultures of hippocampal neurons (Figure 3A), where synapses develop according to a stereotypical sequence of events, from a stage preceding synapse formation (1-3 Days In Vitro; DIV) to a synaptically active mature phase (13-14 DIV) (Matteoli et al, 1995). After the in vitro development, mature cultured neurons obtained from IL-6-exposed embryos exhibited higher mEPSCs frequency, without changes in amplitude, if compared to vehicle-exposed cultures (Figure 3B), whereas mIPSCs were unaffected (Figure 3C). Thus, the effect of IL-6 is intrinsically sculpted in neurons, even after their isolation from the brain context.

**Figure 3.**
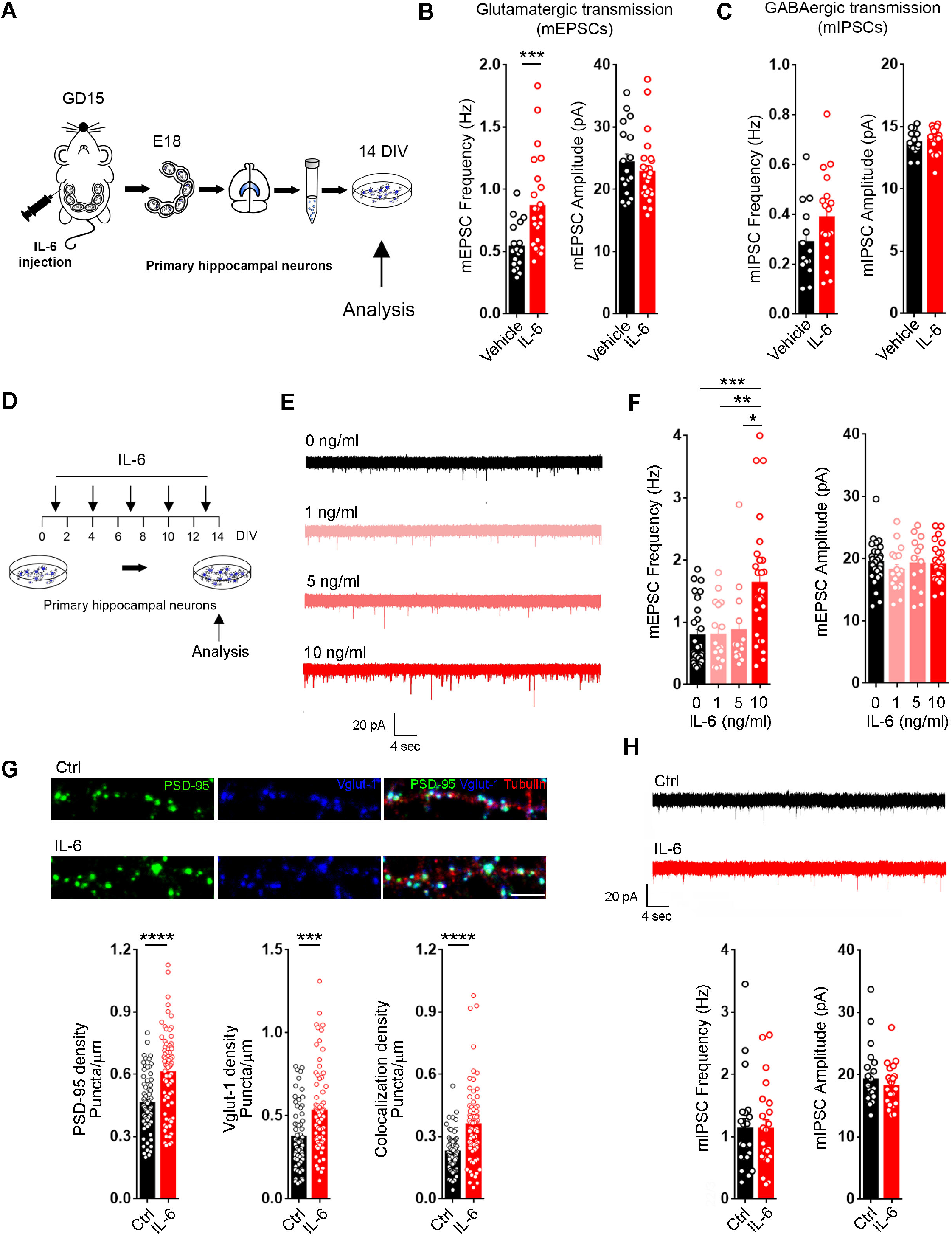
The proinflammatory cytokine IL-6 selectively increases Glutamatergic synaptogenesis in developing hippocampal cultures. **(A)** Representative scheme of the experimental procedure: a single pulse of either vehicle (saline, as control) or IL-6 (5μg) was intraperitoneally injected in pregnant mother at gestational day 15 and primary hippocampal neurons were established at E18. Cultured neurons were assayed at 14 DIV through patchclamp recording. **(B)** Quantitative analysis of mEPSCs frequency and amplitude in cultures established from either vehicle or IL-6 exposed embryos. (Vehicle n = 18 cells, IL-6 n= 25 cells. Three independent experiments). Mann-Whitney test. ***p=0,0005. (**C)** Quantitative analysis of mIPSCs frequency and amplitude in control and upon IL-6 treatment cultures (Vehicle n = 14 cells, IL-6 n= 18 cells. Three independent experiments). Mann-Whitney test. (**D)** Schematic representation of the experimental procedure. Different concentration of IL-6 were incubated throughout the in vitro development of hippocampal neurons, from 1 DIV up to 13 DIV, adding the cytokine every 3 days, and subsequently analyzed at 14 DIV. **(E)** Representative traces of electrophysiological recordings of mEPSC performed in neuronal cultures at 14 days in vitro (DIV) upon chronic treatment with IL-6 at the indicated concentrations. **(F)** Quantitative analysis of both frequency (left panel) and amplitude (right panel) of mEPSCs recorded at different concentration of IL-6. (Cells: Ctrl n=27, 1ng/ml n=17, 5ng n=14, 10ng n=28. Three independent experiments). One-way ANOVA on ranks followed by Dunn’s multiple comparison test ***p=0,0005, **p=0,0063, *p=0,0377. **(G)** Immunofluorescence analyses of glutamatergic synaptic contacts recognized using antibodies against post-synaptic (PSD 95, green) and presynaptic (Vglut1, blue) markers localized along dendritic processes (Beta III Tubulin, red) in control and after chronic treatment with IL-6 10ng/ml. Scale Bar 10 μm. Lower panels: bar graphs showing post-synaptic (left graph) pre-synaptic (central panel) and the colocalizing puncta density (right panel). (dendrites: ctrl n= 64; IL-6 n=70. Four independent experiments). Mann-Whitney test. ****p < 0.0001, ***p=0.0002, ****p < 0.0001. (**H)** Representative traces of electrophysiological recordings of mIPSC performed in neuronal cultures at 14 DIV upon chronic treatment with IL-6 at 10 ng/ml. Lower panel: quantitative analysis of amplitude and frequency of mIPSCs (Cells: Ctrl n=22, IL-6 n=23. Three independent experiments). Mann-Whitney test.

To assess whether IL-6 was effective in promoting synapse formation by acting specifically on neuronal cells, we used primary cultures of embryonic hippocampal neurons directly exposed to IL-6. Hippocampal cultures were incubated with different concentrations of IL-6 (1, 5 and 10 ng/ml) starting from 1 up to 14 DIV, refreshing IL-6 every 3 days (Figure 3D), and the mEPSCs were evaluated at 14 DIV by patch clamp recordings (Figure 3E). The frequency of mEPSCs was significantly increased upon incubation with IL-6 at 10 ng/ml, whereas mEPSCs amplitude was unchanged (Figure 3F). Passive membrane properties, including resting potential (Figure 3SA) and input resistance (Figure 3SB) were not altered by IL-6 treatment at 10ng/ml, thus indicating that the overall health state of neurons was not affected by the cytokine at this concentration. Henceforth, IL-6 was used at 10 ng/ml in cultured neurons. Given the presence of glial cells in cultures, to rule out the possible contribution of astrocytes to the enhanced excitatory neurotransmission, cultured neurons were grown in the presence of the anti-proliferative agent cytosine arabinoside (Ara-C) to obtain pure neuronal cultures (Figure 4SA). The increased frequency of mEPSCs observed upon chronic IL-6 treatment (Figure 4SB,C), demonstrated that astrocytes are not involved in the enhancement of excitatory neurotransmission. It is worth noting that, similar to what obtained in vivo, mIPSCs were not affected by the same IL-6 concentration effective in promoting glutamatergic synapses (Figure 3H), which is indicative of an excitatory/inhibitory (E/I) imbalance due to excessive excitatory inputs also in vitro.

**Figure 4.**
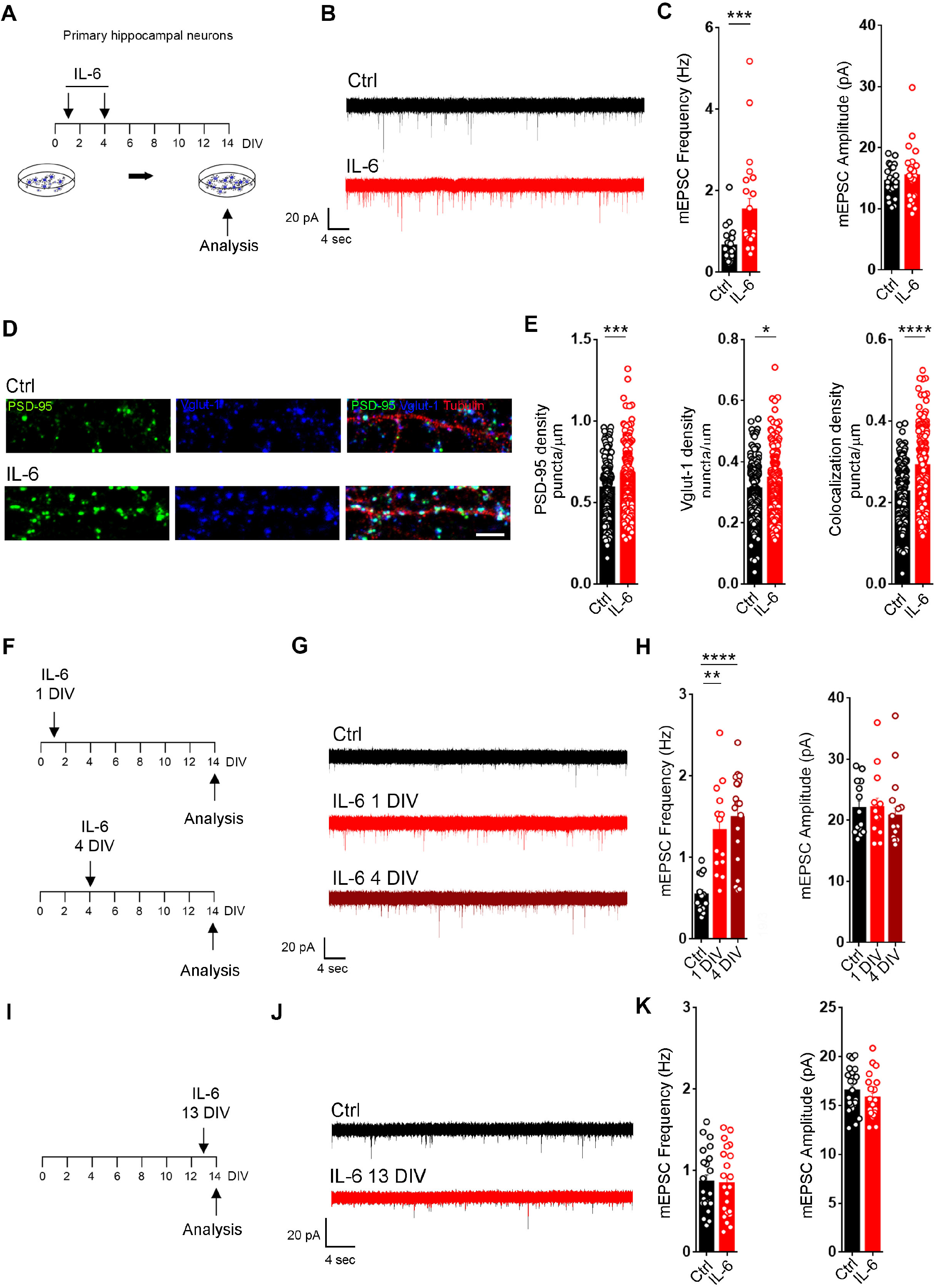
A single IL-6 treatment at early stage of neuronal development is sufficient to increase Glutamatergic synapse at mature stages. **(A)** Schematic representation of the experimental procedure: IL-6 at 10ng/ml was added at early stages of neuronal development at 1 and 4 DIV, and at 7 DIV the medium was replaced with fresh one. Neurons were assayed at 14 DIV. (**B)** Electrophysiological traces of mEPSCs recorded in neuronal cultures at 14 DIV in control and upon a short (1-4 DIV) treatment with IL-6. (**C)** Quantitative analysis of frequency and amplitude of mEPSCs in control condition and upon IL-6. (Cells: ctrl n=24, IL-6 n= 22. Three independent experiments). Mann-Whitney test. ***p=0,0007. **(D)** Immunofluorescence analysis of glutamatergic synaptic density identified using pre-synaptic (Vglut-1, in blue) and post-synaptic (PSD-95, in green) markers along a dendritic branch stained with beta III tubulin, (in red). Scale bar, 10 μm. **(E)** Quantitative analysis of PSD-95 puncta density (left panel), Vglut-1 puncta density (middle panel) and colocalizing puncta density (right panel) in both conditions. (Dendrites: ctrl n= 133, IL-6 153. Three independent experiments) Student’s t-test. ***p=0,0002, *=p<0,0126, ****p<0,0001. **(F)** Schematic representation of the experimental procedure: a single exposure of IL-6 10 ng/ml was performed at 1 DIV (Upper panel) or at 4 DIV (lower panel), and neurons were assayed at 14 DIV. **(G)** Representative traces of mEPSCs recorded at 14 DIV in control condition and upon single IL-6 exposures at the indicated time points. **(H)** Quantitative analysis of both frequency and amplitude of mEPSC at 14 DIV in the two conditions. (Cells: Ctrl n= 14, IL6 1DIV n=14, IL6 4DIV n=19. Three independent experiments.). One-way ANOVA on ranks followed by Dunn’s multiple comparison test. **p=0,0011; ****p<0,001. (**I)** Schematic representation of the experimental procedure: a single exposure of IL-6 10 ng/ml was performed at 13 DIV, and neurons were assayed at 14 DIV. (**J)** Representative traces of mEPSCs recorded at 14 DIV in control condition and upon single IL-6 exposures at later stage of development. (**K)** Quantitative analysis of both frequency and amplitude of mEPSC in control and upon IL-6 treatment at later developmental stages. (Cells Ctrl n=22, IL-6 n=22. Three independent experiments) Mann-Whitney test.

To directly address whether the increased glutamatergic basal transmission results from a higher number of excitatory synapses, glutamatergic synaptic density was investigated in both control and treated cultures by using antibodies against the presynaptic vesicular glutamate transporter-1 (V-glut-1) and postsynaptic density-95 (PSD-95) (Figure 3G, upper panel). IL-6-treated cultures displayed a significantly increased density of Vglut-1 and PSD-95 positive puncta per unit length of parent dendrite, as well as higher extent of colocalization between Vglut-1 and PSD-95, indicating a higher number of structurally mature synapses along the dendritic branches (Figure 3G, lower panels).

The increase in mEPSCs frequency might in principle result from different mechanisms, including enhanced release probability at presynaptic terminals. To exclude this possibility, short term plasticity was evaluated through paired recording measurements between synaptically connected neurons (Figure 4SD) (Maximov et al, 2007). Untreated neurons exhibited a paired pulse facilitation (PPF) at different interpulse intervals (Figure 4SE), with the paired pulse ratio progressively reducing as the interpulse intervals increased, as already described (Farisello et al, 2013; Nanou et al, 2016). Such trend was not significantly altered in neurons subjected to IL-6 chronic treatment, thus excluding that IL-6 enhances presynaptic release probability (Figure S4E). Notably, the amplitude of post synaptic currents (EPSCs) evoked by a single action potential was significantly higher in neurons exposed to IL-6 treatment (Figure S4F), in line with the increase in glutamatergic inputs (Chao et al, 2007).

We also ruled out that the increase in glutamatergic synapses resulted from homeostatic compensatory mechanisms (e.g. synaptic scaling), as a consequence of reduced neuronal excitability upon IL-6 treatment (Vereyken et al, 2007; Wierenga et al, 2006). Indeed, electrically-evoked calcium transients recorded through single cell calcium imaging (Figure S4G,H) (Bedogni et al, 2016; Pozzi et al, 2013) showed a comparable neuronal excitability in control and IL6-treated cultures (Figure S4I). Finally, the lack of changes in the total number of cells, as well as in the ratio between astrocytes and neurons, in cultures chronically exposed to IL-6 (Figure S4J-L) excluded a role for the cytokine in modulating the overall cellular components of the culture.

### A single pulse of IL-6 at stages preceding synaptogenesis is sufficient to promote a long-lasting increase of glutamatergic synaptic transmission

The chronic treatment with IL-6 in cultured neurons does not perfectly reflect the transient maternal IL-6 elevation induced in vivo, where embryos were exposed to a single pulse of IL-6 (see Figure 1). Hence, to probe whether a shorter incubation with IL-6 was still effective in promoting glutamatergic transmission, neuronal cultures were incubated with the cytokine at restricted phases of early neuronal development (Figure 4A). IL-6 treatment performed at 1 and 4 DIV, was sufficient to increase mEPSC frequency (Figure 4B,C) and glutamatergic synapse density at 14 DIV (Figure 4D,E). Furthermore, we found that even a single pulse of IL-6, applied either before (1DIV) or during (4 DIV) the synaptogenesis process (Figure 4F), was sufficient to boost glutamatergic synaptic transmission (Figure 4G,H). Conversely, a single pulse of IL-6 at synaptically active mature stages (13 DIV) (Figure 4I) failed to increase glutamatergic transmission (Figure 4J,K). These results indicate that IL-6 does not affect synaptic transmission acutely and that a single, transient IL-6 elevation at stages preceding synapse formation promotes a long-lasting selective enhancement of glutamatergic synapses. To test the specificity of IL-6 to act as prosynaptogenic molecule, we extended our analysis to other proinflammatory cytokines, such as INFγ, TNFα and IL1β, known to play a central role in inflammation. The cytokines were individually applied to neuronal cultures at 1 DIV and glutamatergic transmission was assessed at mature stages (Figure S5A). Unlike to IL-6, none of the proinflammatory cytokines tested was effective in enhancing excitatory transmission (Figure S5B), indicating specific action of IL-6 as prosynaptogenic molecule.

### STAT3 activity is required for the IL-6 dependent increase of glutamatergic synapses

To gain insights into the molecular underpinnings of the IL-6-mediated effect, we explored IL6-dependent signaling pathways involved in this process. Given the long-lasting effect produced by IL-6 on glutamatergic synapses, it is plausible that a transcriptional-dependent mechanism might be engaged. IL-6 is known to activate a cascade of molecular events converging on the activation of the transcription factor Signal Transducer and Activator of Transcription 3, STAT3 (Heinrich et al, 2003). This prompted us to investigate the involvement of STAT3 in our process. We found that both transcript and protein levels of STAT3 were enhanced in neuronal cultures upon short IL-6 treatment (Figure 5A-D). To note, IL6-mediated upregulation of STAT3 occurred also in AraC-treated cultures, indicating a neuronal-specific STAT3 activation (Figure 5C, D). Accordingly, STAT3 upregulation was tested in the embryonic hippocampi after 24 h from IL-6 i.p. injection (Figure 5E), thus confirming its modulation in vivo (Figure 5F,G).

**Figure 5:**
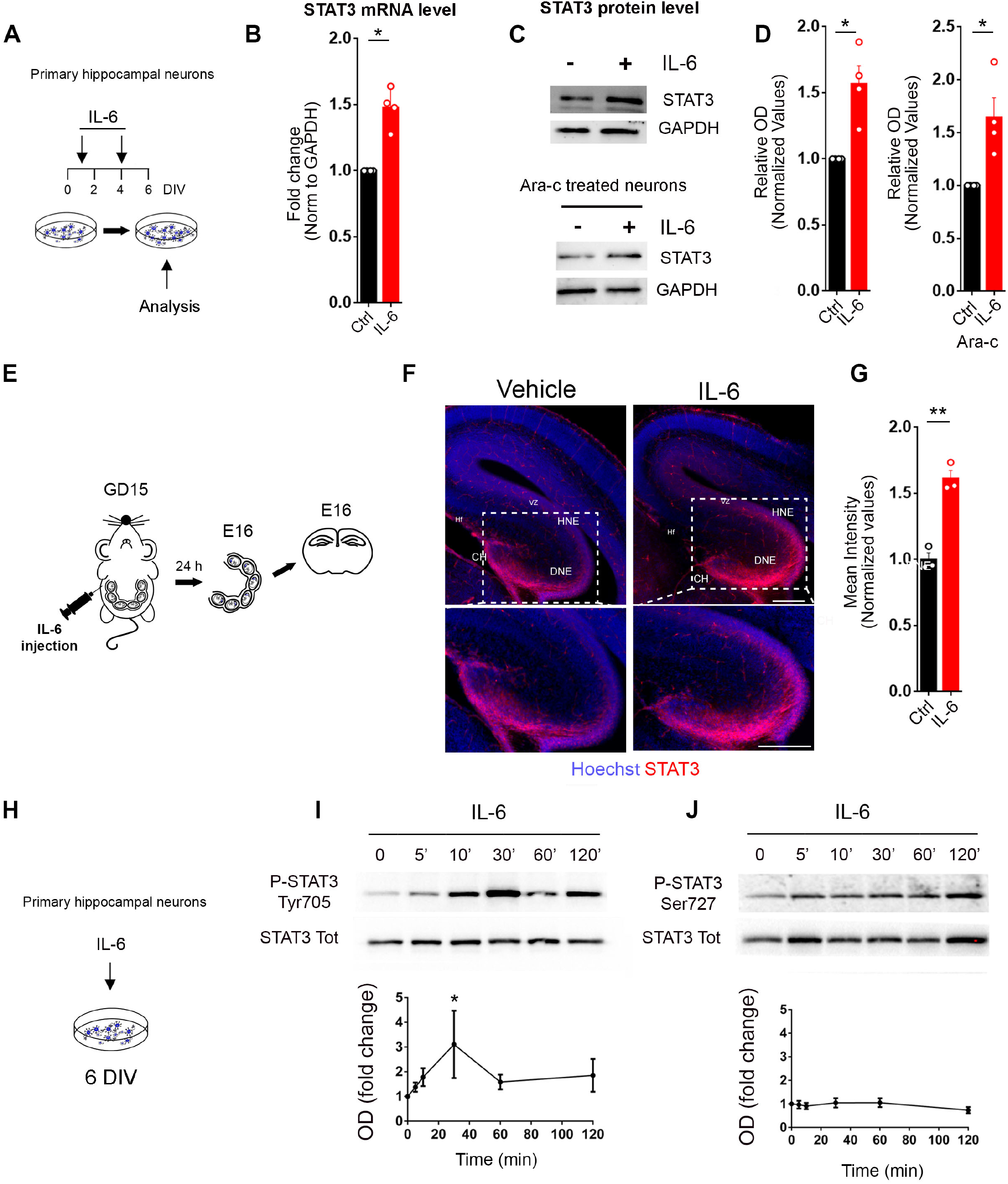
IL-6 increases STAT3 levels both in vitro and in vivo in neurons and induces a selective phosphorylation at tyrosine 705. **(A)** Schematic representation of the experimental procedure: IL-6 at 10ng/ml was added at 1 and 4 DIV, and at 6 DIV neurons were collected for the analysis. (**B)** Quantitative qPCR of STAT3 mRNA levels in control and IL-6 treated cultures (four independent experiments). One sample t test. *p=0.008. (**C)** Western blot analysis of STAT3 protein levels in control and IL-6 treated neuronal cultures both in absence (with astrocytes, upper panel) or in the presence of Ara-C (no astrocytes, lower panel). (**D**) Quantitative analysis of the optical density of STAT3 immunoreactive bands normalized by GAPDH level, in normal (left graphs) and in Ara-C treated cultures (right graphs). (Four independent experiments). One sample t test. Ctrl cultures:*p=0.0254; Ara-C cultures: *p=0.04. (**E)** Representative scheme showing the experimental procedure: a single pulse of either vehicle (saline, as control) or IL-6 (5μg) was intraperitoneally injected in pregnant mothers at embryonic day 15 and the hippocampus of embryos were analyzed after 24 hours from the injection through immunofluorescence. (**F)** Representative immunofluorescence of hippocampal regions stained with STAT3 (red) and Hoechst (blue), established from embryos at E16 exposed to vehicle or IL-6 at E15 via maternal intraperitoneal injection. (VZ: ventricular zone, HNE: hippocampal neuroepithelium, DNE: dentate neuroepithelium, CH: cortical hem) scale bar 250μm. Inset panels: higher magnification of the selected area. Scale bar: 250 μm. (**G)** Quantitative analysis of STAT3 mean intensity at hippocampal level in the two conditions (Vehicle n= 3 embryos, IL-6 n= 3 embryos) Student’s t-test. Data shown ad mean± SEM. **p=0,0098. (**H)** Scheme of the experimental procedure: cultured neurons at 6 DIV were treated acutely with IL-6 and collected at different time points for western blot analysis. (**I-J)** Time course analysis of STAT3 phosphorylation at tyrosine 705 and **(J)** serine 727 upon acute (30 min) IL-6 treatment of 6 DIV cultured neurons. Quantitative analysis of the optical density of STAT3 phosphorylation levels in tyrosine 705 (I, lower panel) and serine 727 (J, lower panel) normalized against the total amount of STAT3 (Four independent experiments). Wilcoxon Signed Rank Test. *p=0.0313

STAT3 is known to undergo phosphorylation at two specific residues, Tyrosine-705 (Tyr-705) and Serine-727 (Ser-727). Whilst the phosphorylation of Tyr-705 is known to be IL-6-dependent (Wen et al, 1995), the phosphorylation at Ser-727 varies in different cell types according to the type of stimulation triggered (Chung et al, 1997a; Chung et al, 1997b; Lim & Cao, 1999). Analysis of STAT3 phosphorylation after an acute IL-6 application in hippocampal neurons (Figure 5H) revealed a transient increase of phosphorylation at Tyr-705 (Figure 5I), whilst Ser-727 remained unaltered (Figure 5J), indicating that IL-6 directly engages STAT3 activation.

To next assess the contribution of STAT3 activation on glutamatergic synaptogenesis, we took advantage of a pharmacological compound, Stattic, which selectively prevents STAT3 phosphorylation (Schust et al, 2006). Cultured neurons were transiently exposed to a single pulse of IL-6 (at 1 DIV, see scheme in Figure 4F) in the presence of either vehicle (as control) or 1 μM Stattic. This concentration was effective in blocking STAT3 phosphorylation (Figure 6A,B), without affecting neuronal survival (Figure S5C) nor synaptic basal transmission (Figure S5D). Patch clamp analysis of glutamatergic basal transmission (Figure 6C) showed that the IL-6-dependent increase of mEPSC frequency (Figure 6D) was prevented by Stattic. Accordingly, the higher glutamatergic synaptic density (Figure 6E,F) induced by IL-6 does not occur in the presence of STAT3 inhibitor. Noteworthy, Stattic also prevented the increase of glutamatergic transmission induced by chronic treatment with IL-6 (Figure S6A-C), without affecting the overall increase of STAT3 protein level (Figure S6D), indicating that the IL-6-mediated upregulation of the transcription factor is not self-sustained by STAT3 itself. Moreover, IL-6 treatment did not change the expression of a panel of pre-post-synaptic proteins, nor the expression of the transcription factor NFKβ, involved in many immune-dependent processes (Figure S6E). Furthermore, the short application of IL-6, effective in enhancing excitatory synaptic transmission (Figure 3), did not result in a sustained upregulation of STAT3 protein at later stages (Figure 6SF-G), indicating that the transcription factor is only transiently induced by IL-6.

**Figure 6:**
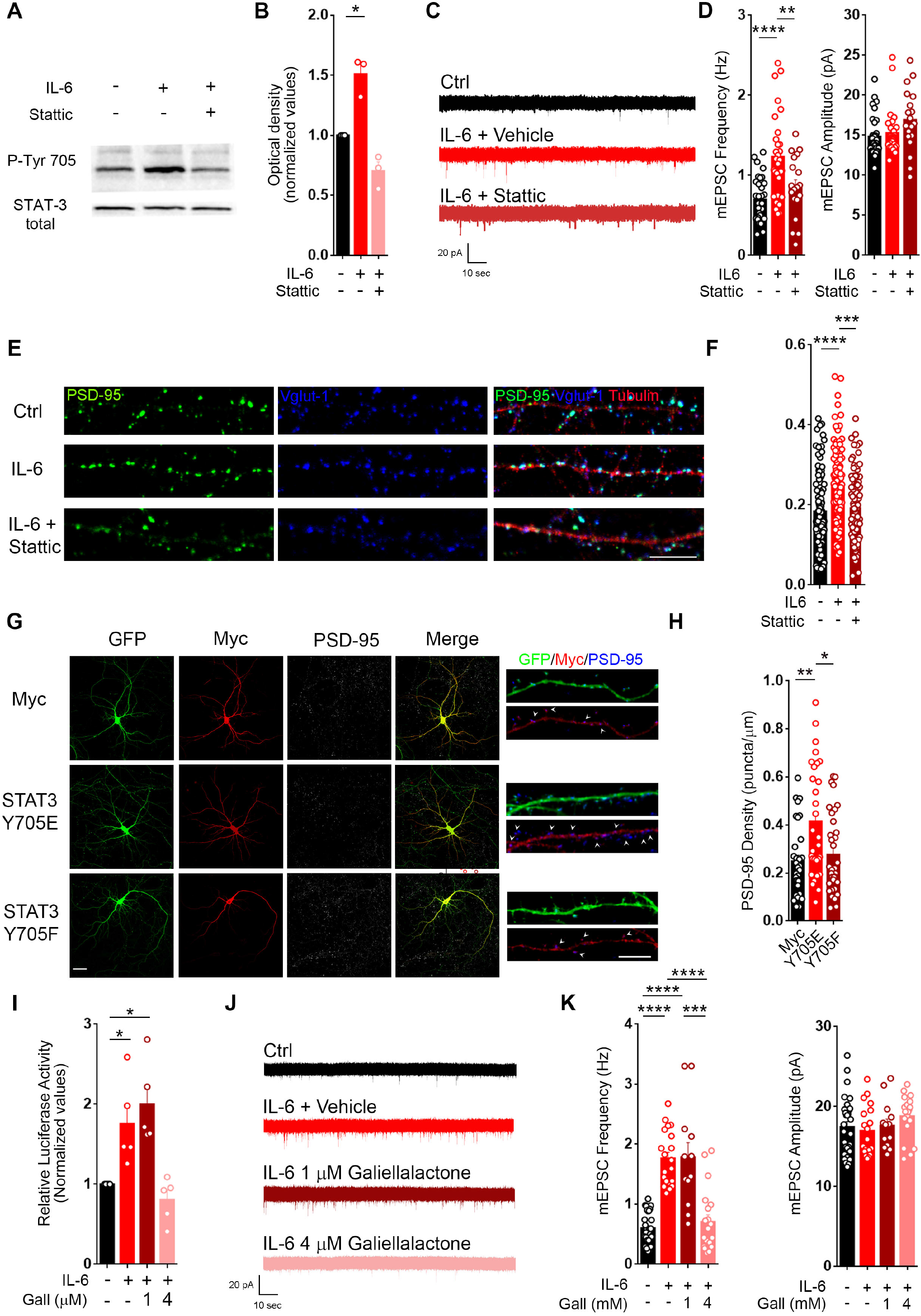
STAT3 genomic activity is causally linked with the IL6-dependent increase of glutamatergic synapses. **(A)** Western Blot analysis of STAT3 phosphorylation in cultured neurons at 6 DIV acutely treated (30 min) with IL-6 in absence or presence of Stattic 1μM **(B)** Mean optical density quantification of STAT3 phosphorylation normalized on total STAT3 protein level. (three independent experiments). One sample t test. *p=0,0326. **(C)** Representative electrophysiological traces of mEPSCs recorded in cultured neurons at 14 DIV in control condition and upon a single application of IL-6 at 1 DIV (see scheme in figure 2F) in the presence of either vehicle or 1μM Stattic. (**D)** Quantitative analysis of mEPSCs frequency and amplitude in the different conditions (Ctrl n=33 cells, IL-6 n=30 cells, IL-6 stattic n= 19 cells. Three independent experiments.) One-way ANOVA followed by Tukey’s multiple comparisons test. ****=p<0,001; **=p<0,0065. **(E)** Immunofluorescence analysis of glutamatergic synaptic density evaluated in the same conditions through antibodies against pre-(Vglut1, blue) and post (PSD-95, red) synaptic markers along dendritic branches (beta III tubulin, in red). Scale bar 10 μm. **(F)** Quantitative analysis of colocalizing puncta density in the different conditions. (Dendrites: Ctrl n=117, IL-6 n=130, IL-6 + stattic n= 88). One-way ANOVA on ranks followed by Dunn’s multiple comparison test. ***p < 0.0004; ****p < 0.0001. **(G)** Representative images of hippocampal neurons expressing GFP together with: myc alone (upper panels), myc-flagged STAT3–phosphomutant Y705E (mimicking a constitutive phosphorylated state) (middle panels), myc-flagged STAT3–phosphomutant Y705F (mimicking a constitutive non-phosphorylated state) (lower panels). These neurons were stained with antibodies against myc (red), to visualize the mutant expression, and with PSD-95 (blue), to evaluate the density of post-synaptic puncta along the GFP-expressing (green) dendritic branches. High-magnification images on the right, arrows indicated PSD-95 positive puncta). Scale bar left panel 20μm, right panel 10 μm. **(H)** Quantitative analysis of PSD-95 positive puncta density and size in the three different conditions respect to control condition. (dendrites: Y705E=33; Y705F=30; MYC=35. Three independent experiments) One-way ANOVA on ranks followed by Dunn’s multiple comparison test. *p=0,0258; **p=0,0057. (**I)** STAT3 luciferase reporter assay performed in cultured neurons stimulated with IL-6 for 48 hours alone and in the presence of Galiellalactone 1 and 4 μM. (Five independent experiments) One sample t test. IL-6 *p=0,0353; IL-6 Gall*p=0,0122. **(J)** Representative electrophysiological traces of mEPSCs recorded in cultured neurons at 14 DIV in control condition and upon a single application of IL-6 at 1 DIV with either vehicle or Galiellalactone at 1 and 4 μM. **(K)** Quantitative analysis of mEPSCs frequency and amplitude in the indicated conditions. (Cells: Ctrl n=30, IL-6 n=20, IL-6 Gall1μM n=13, IL-6 Gall4μM n=20. Three independent experiments) One-way ANOVA on ranks followed by Dunn’s multiple comparison test ***p=0,0008; ****p<0,0001.

To univocally demonstrate that STAT3 activation *per se* is sufficient to boost glutamatergic synapses in a neuron-autonomous fashion, we took advantage of two STAT3 mutant forms: the mutant Y705F, in which the tyrosine 705 residue is replaced by phenylalanine, thus mimicking a constitutively unphosphorylated state of the protein (inactive form), and the mutant Y705E, in which the tyrosine 705 residue is replaced by a glutamate, thus mimicking a constitutively phosphorylated state (active form). Both STAT3 phosphomutants were translated as fusion-proteins bearing a Myc-tag, and the synaptic density analysis was performed in sparse transfected neurons expressing the fusion proteins (Figure 6G). Analysis of PSD-95 positive puncta density in the transfected neurons showed that the phosphomimetic mutant Y705E, but not the Y705F, significantly increases both PSD-95 density and size (Figure 6H). These data indicate that STAT3 transient activation is required for the IL-6 effect on glutamatergic synapses.

### IL-6 promotes glutamatergic synaptogenesis through a STAT3 dependent genomic effect involving RGS4 activity

As a transcription factor, STAT3 is known to modify many cellular processes through genomic effects (Heinrich et al, 1998). However, a recent evidence reported a non-genomic effect of STAT3 in modulating synaptic plasticity (Nicolas et al, 2012). To assess whether STAT3 activation affects glutamatergic transmission via genomic or non-genomic mechanisms, the fungal metabolite Galiellalactone, a selective STAT3 inhibitor able to prevent STAT3 binding to DNA without affecting its phosphorylation (Weidler et al, 2000), was employed. The compound working concentration was selected through a luciferase reporter assay probing STAT3 genomic activity. 4 μM Galiellalactone was identified as the effective concentration able to block STAT3 genomic activity (Figure 6I) interfering with neither synaptic basal transmission (Figure S7D) nor the IL-6 dependent STAT3 phosphorylation (Figure S7C), and with no effect on neuronal survival (Figure S7A,B). Interestingly, this concentration was able to block the increase of mEPSC frequency upon IL-6 (Figure 6 J,K), thus confirming that the pro-synaptogenic activity of IL-6 is mediated by the genomic effect induced by STAT3 activation.

In order to identify the molecular pathways involved in IL6-induced glutamatergic synaptogenesis, we investigated the overall transcriptional rearrangement induced by IL-6 through STAT3 activation by 10X single cell transcriptomics (scRNA sequencing). We assessed the genome-wide expression profiling of 3,687 single cells isolated from cultured hippocampal neurons in control conditions or upon IL-6 application (ctrl: 1865; IL-6: 1822) at DIV5 (Figure 7A). The median number of genes detected per cell was 5,229 for IL6 and 4,714 for ctrl, and the median number of transcript (unique molecular identifiers, UMI) per cell was 22,011 and 17,572 respectively. Unsupervised clustering analysis was applied on our pooled dataset using Seurat R package (Butler et al, 2018) at various resolution to computationally define cellular heterogeneity. The resolution tree of the clustering is reported in Figure S8A. The optimal cluster resolution (0.6) - selected based on cluster index stability and number of identified clusters - resulted in eight transcriptionally distinct populations were identified according to their gene transcriptional profile (Figure 7B-C). The clusters were represented in the two experimental conditions and substantially overlapped (Figure 7D), suggesting that changes in the transcriptional profiling within each single cluster, rather than modifications of the distinct cell identities, occurred upon IL6 treatment.

**Figure 7:**
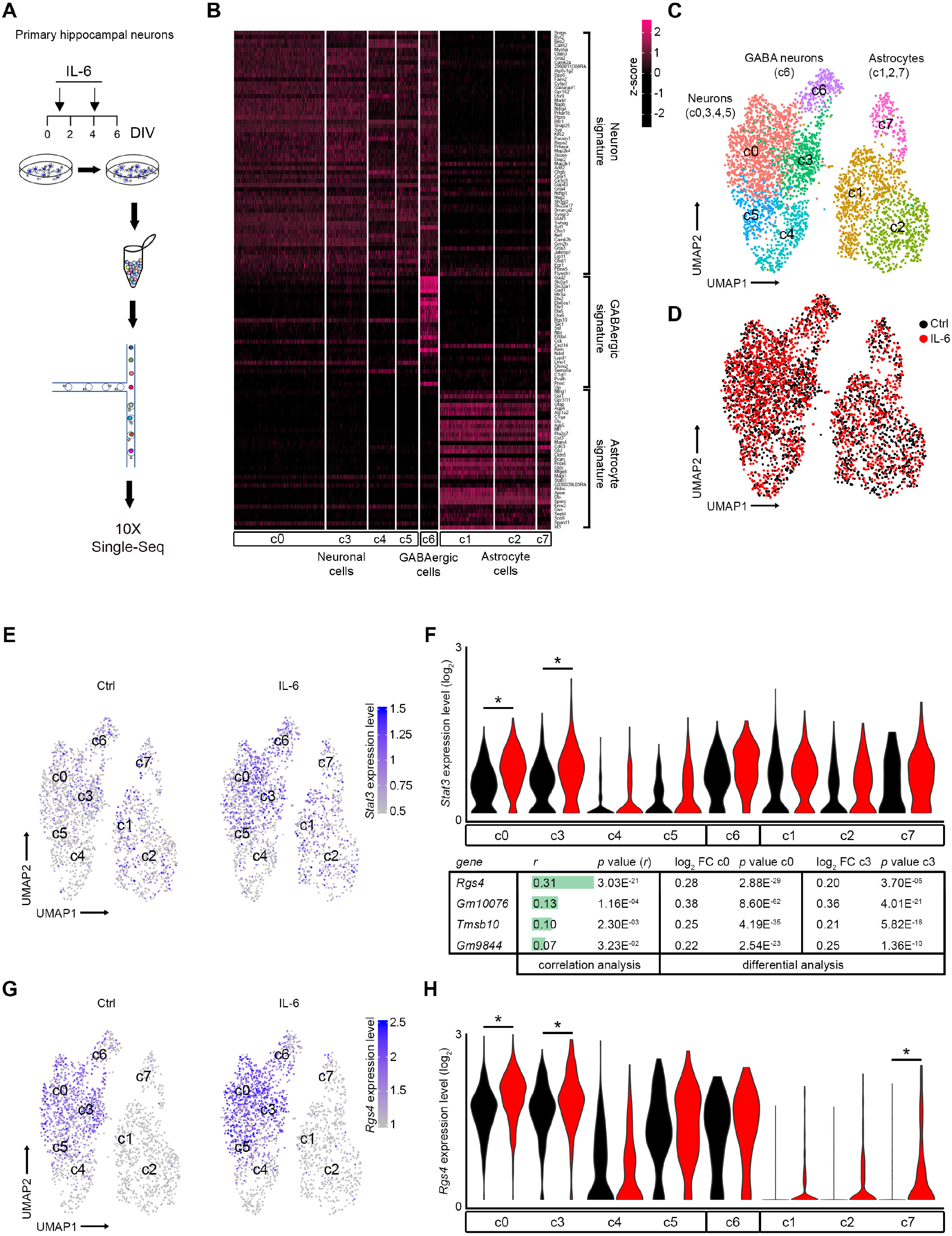
Single cell transcriptional analysis of developing neurons reveals the upregulation of *RGS4* gene in specific neuronal cluster cells upon IL-6 treatment. **(A)** Schematic representation of the experimental workflow. Cultured neurons were incubated with IL-6 for a short period (1 and 4 DIV) and collected at 5 DIV for single cell sequencing analysis. (**B)** Heat map showing the average expression value of Neuron and Astrocyte gene signatures (Lein et al, 2007) and manually-curated GABAergic signature across all cells belonging to cluster 0 to 7. Average expression scale is shown on the right. **(C)** UMAP (uniform manifold approximation and projection) projection of all detected single cells. Each point depicts a single cell, colored according to cluster designation. n = 3,687 individual cells. Clusters are labeled according to cell enrichment assessed by AUCell R Bioconductor package (see Supplementary Figure S8). **(D)** UMAP projection of all detected single cells, colored according to experimental condition. Black points: control cells, red points: IL-6 treated cells. **(E)** UMAP projection of all detected single cells, colored according to *Stat3* gene expression level. Average expression scale is shown on the right. Left panel: control cells, right panel: IL-6 treated cells. **(F)** Top: Expression distribution (violin plots) showing *log*-transformed, normalized expression levels of *Stat3* gene, in all clusters grouped by cell type and coloured according to experimental condition. Black points: control cells, red points: IL-6 treated cells. *: *p* value ≤0.05, Wilcoxon Rank Sum test. Bottom: Table showing Pearson *r* correlation value of genes significantly correlated with *Stat3* gene expression level in cluster 0 and cluster 3 of IL-6 treated cells, and significantly modulated after IL-6 treatment in cluster 0 and cluster 3. **(G)** UMAP projection of all detected single cells, colored according to *Rgs4* gene expression level. Average expression scale is shown on the right. Left panel: control cells, right panel: IL-6 treated cells. **(H)** Expression distribution (violin plots) showing *log*-transformed, normalized expression levels of *Rgs4* gene, in all clusters grouped by cell type and colored according to experimental condition. Black points: control cells, red points: IL-6 treated cells. *: *p* value ≤0.05, Wilcoxon Rank Sum test.

We, then, systematically classified the cells by comparing their transcriptional profiles to pre-existing signatures gene sets of endogenous neuronal and glial types (Cahoy et al, 2008; Cembrowski et al, 2016; Harris et al, 2018; Lein et al, 2007) (Supplementary Table 1), see materials and methods for details), to automatically evaluate their respective gene enrichment. We thus defined eight main transcriptionally distinct cell types reflecting the diversity of neuronal and non-neuronal classes found in the hippocampus (Cembrowski et al, 2016; Pelkey et al, 2017).

Based on this enrichment analysis, we found that three out of the eight clusters belonged to the astrocyte lineage (c2,7,8), expressing glial specific marker gene including *Gfap* and *Aqp4* with one of them showing markers of cycling cells (c2) (Figure S8E and Supplementary Table 1). The clusters c0,c3,c4,c5 and c6 display enrichment, instead, of neuronal-specific genes, among others, *Snap25, Syp* and *Syt1*, which are expressed in hippocampal neurons. Among these neuronal clusters, only one could be clearly ascribed to the GABAergic lineage (c6), expressing markers prototypical of cortical and hippocampal GABAergic neurons including *GAD1* and *Slc32a1*(Figure S8B-D, and Supplementary Table 1) (Mancinelli & Lodato, 2018). In terms of relative abundance of distinct cellular populations, the single cell sequencing data also provided evidence that our in vitro culture system support the development of the different classes, while, at large, respecting the ratio of excitatory and inhibitory neurons, and glial cells, which is appropriate for the developmental stage of the analysis (Lodato & Arlotta, 2015; Mancinelli & Lodato, 2018; Pelkey et al, 2017)

To further investigate the transcriptional changes induced by transient IL-6 application, we performed differential expression analysis between IL-6-stimulated conditions compared to control. We identified 63 differentially expressed genes (Adjusted p-value cut-off of 0.05 and a logfc.threshold=0.20), with 55 being upregulated and 8 being downregulated upon IL6 treatment. Notably, as expected *Stat3* gene expression was significantly upregulated (p value ≤0.05), but only in the neuronal clusters (c0,c3), indicating a predominant activation of STAT3 in specific neuronal cell types upon IL-6 stimulation. Noteworthy, IL-6-mediated *Stat3* expression was not subjected to any significant modifications in either astrocyte or GABAergic neuron cluster (Figure 7E-F).

In order to identify possible *Stat3 co-*associated genes, we then combined an unbiased correlation analysis within neuronal clusters where *Stat3* was found significantly upregulated (c0;c3), together with a differential analysis between IL-6-stimulated cultures and controls. Only four genes show significant correlation with *Stat3* expression upon IL-6 treatment (Figure 7F, bottom table). Among them, the highest correlation (positive) was found between S*tat3* and *Rgs4* (r = 0.31), indicating that in those clusters where *Stat3* was upregulated, *Rgs4* expression was concomitantly increased (Figure 7 G,H), ranking at the top of significantly correlated genes. Interestingly, this gene was highly enriched in neuronal clusters (Figure 7H). Furthermore, when we analyzed the promoter region of *Rgs4* through bioinformatics tools for transcription factor binding sites prediction (Khan et al, 2018; Sandelin et al, 2004), we found distinct putative sites containing Stat3 response elements predicted with relative profile score threshold 80% (Figure S8F). These results suggest that *Rgs4* could represent a downstream target gene of STAT3 upon IL-6 stimulation.

The data obtained through single-cell sequencing approach were then validated through qPCR quantitation of *Rgs4* mRNA levels in primary cultured neurons upon acute IL-6 treatment. *Rgs4* was significantly upregulated upon IL-6 treatment in a STAT3 dependent mechanism, as Galiellalactone was able to completely prevent the IL-6 mediated mRNA elevation (Figure 8A). STAT3 activity induced RGS4 transcriptional expression in a cell autonomous fashion. This was demonstrated by transfecting mouse N2A neuronal cell lines with different plasmids expressing either the active (STAT3-Y705E) or the inactive (STAT3-Y705F) form of STAT3 together with a plasmid encoding GFP as a report (Figure 8B). qPCR quantification of RGS4 mRNA expression in FACS-sorted GFP^*+*^ cells, revealed that only the active form of STAT3 induces a significant enhancement of RGS4 expression (Figure 8C).

**Figure 8:**
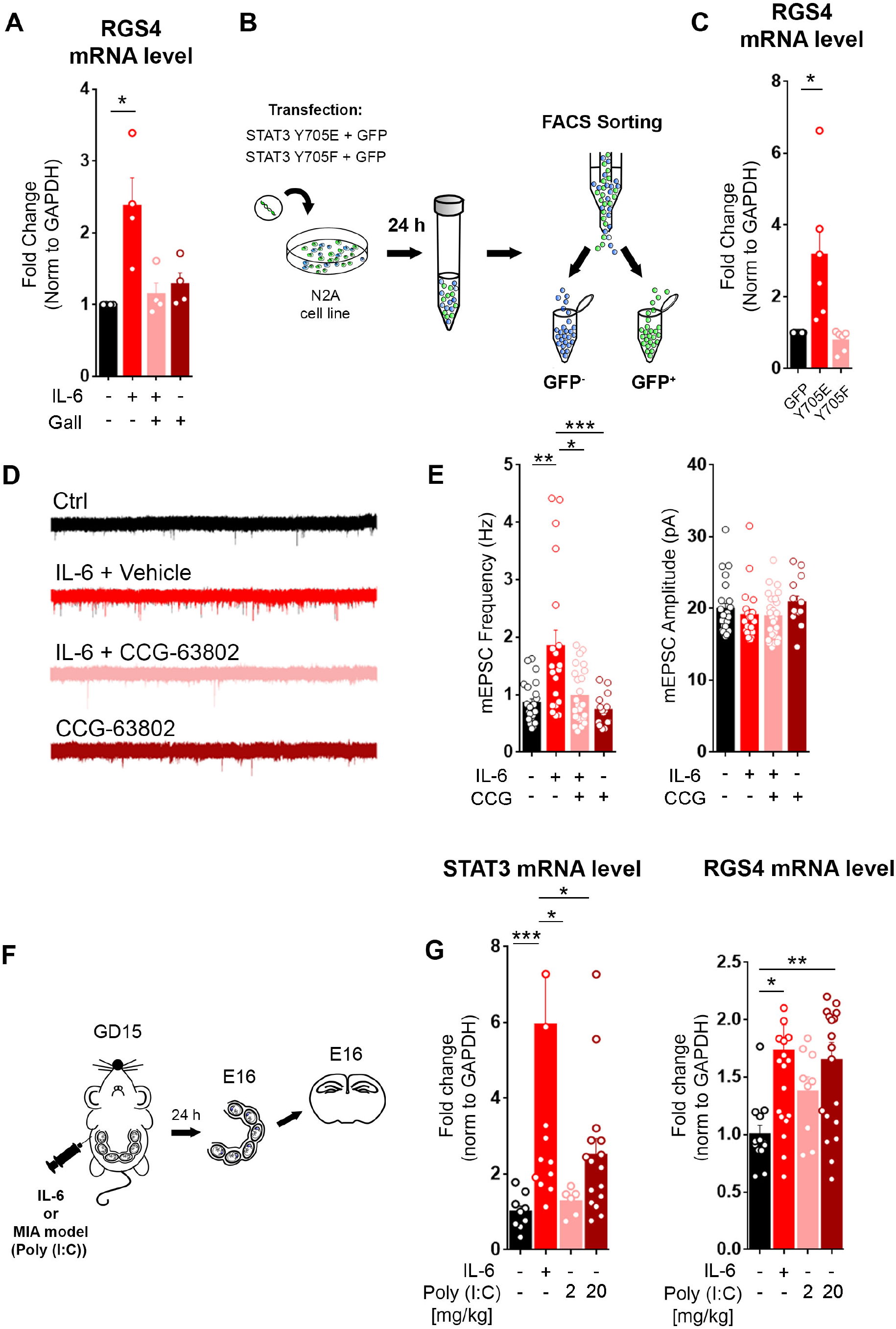
RGS4 activity is required for the IL6-mediated effect on glutamatergic synapses and its upregulation occurs in the brain of embryos upon MIA model. **(A)** qPCR analysis of RGS4 mRNA levels in cultured neurons at 6 DIV upon short IL-6 treatment (1 and 4 DIV as shown in the scheme of Figure 5A) with and without 4 mM Galiellalactone normalized on GAPDH expression. (four independent experiments). One sample t Test *p=0,0389 **(B)** Schematic workflow: N2A cell lines were co-transfected with plasmids coding for the STAT3 phosphomutants Y705E and Y705F, together with GFP. After 24 hours GFP^+^-cells were FACS-sorted and assessed for RGS4 expression level through qPCR analysis. **(C)** Quantitative analysis of RGS4 mRNA level normalized on GAPDH expression in the indicated conditions. (six independent experiments) One sample t test *p=0,0411 (**D)** Representative traces of mEPSCs recorded in cultured neurons at 14 DIV in control condition and upon a single application of IL-6 at 1 DIV (see scheme in figure 2F) in the presence or absence of the RGS4 inhibitor CCG-63802. (**E)** Quantitative analysis of mEPSCs frequency and amplitude in the indicated conditions. (Cells: ctrl n=23, IL-6 n=20, IL-6 CCG n=25, CCG n=15. Three Independent experiments). One-way ANOVA on ranks followed by Dunn’s multiple comparison test **p=0,0046; *p=0,0391; ***p=0,0008. (**F)** Experimental procedure used for MIA model: a single pulse of either vehicle (saline, as control), IL-6 (5μg) or Poly:IC (at 2 or 20mg/Kg) was intraperitoneally injected in pregnant mothers at embryonic day 15 and the hippocampus of embryos were analysed after 24 hours from the injection through qPCR analysis. **(G)** Quantitative analysis of STAT3 and RGS4 expression normalized on GAPDH level in the hippocampi of embryos at the indicated conditions. (Vehicle n= 13 mice, IL-6 n=19 mice, PolyI:C 2mg n= 9 mice, PolyI:C 20mg n=19 mice. Three Independent experiments). One-way ANOVA on ranks followed by Dunn’s multiple comparison test. (STAT3 analysis: ***p=0,0001, *p=0,0125; *p=0,0394. RGS4 analysis: *p=0,0238; **p=0,0074).

To next provide a causal link between RGS4 upregulation and the IL-6-mediated increase of glutamatergic synapses, we took advantage of the recently identified small-molecule CCG-63802, a selective RGS4 inhibitor (Blazer et al, 2010). Neuronal cultures were treated with IL-6 at 1 DIV (see scheme in Figure 4F) in the presence of either the RGS4 inhibitor or vehicle and the excitatory synaptic transmission was monitored at 14 DIV (Figure 8D). The inhibition of RGS4 completely prevented the IL-6-dependent enhancement of glutamatergic transmission (Figure 8E), thus demonstrating that STAT3 −dependent RGS4 elevation is required for the increase of glutamatergic synaptic contacts induced by IL-6.

Finally, to directly investigate the engagement of RGS4 genes in *in vivo* models of prenatal inflammation, pregnant mice were intraperitoneally injected at GD15 with either IL-6 or with Polyinosinic:polycytidylic acid (poly (I:C)), a synthetic analog of viral double-stranded RNA, a well-established model of Maternal Immune activation (MIA) (Choi et al, 2016; Corradini et al, 2017; Hsiao & Patterson, 2011; Shin Yim et al, 2017; Smith et al, 2007a). Poly I:C was used at two different concentrations, 2mg/kg or 20mg/kg, and the expression levels of STAT3 and RGS4 were evaluated after 24 hours from the injection in embryonic hippocampi (Figure 8F). In accordance with the result obtained upon IL-6 injection, the higher Poly I:C concentration was effective in promoting the upregulation of both STAT3 and RGS4 (Figure 8G), thus confirming the involvement of these two genes in MIA models.

## Discussion

Synapse formation represents a fundamental key step in the brain developmental program which may be profoundly affected by genetic defects. However, environmental stressors occurring at this stage, among all inflammatory conditions, may have long-term effects on physiological trajectories, affecting brain connectivity and behavior in adulthood (Boulanger, 2009; Deverman & Patterson, 2009). Although chronic inflammation is recognized to have detrimental consequences on many aspects of neuronal functioning (Glass et al, 2010; Kotas & Medzhitov, 2015; Mandolesi et al, 2015; Najjar et al, 2013; Vezzani et al, 2011), inflammation associated with prenatal infections is likely to be an acute phenomenon (Benedusi et al, 2015; Garetto et al, 2016; Kallikourdis, 2018; Munoz-Suano et al, 2012), with a long-range impact on synaptogenesis which is completely unknown. This aspect has a crucial relevance given the clear association between prenatal immune activation and neurodevelopmental diseases, such as ASD and schizophrenia, occurring during childhood and adolescence (Boulanger-Bertolus et al, 2018; Estes & McAllister, 2016; Knuesel et al, 2014).

In this study, using a combination of in vitro and in vivo models, we demonstrated that a transient elevation of IL-6 levels at early developmental stages is sufficient to exert a specific long-lasting effect on glutamatergic synaptogenesis, resulting in excessive density of excitatory inputs accompanied by enhanced synaptic basal transmission. The pro-synaptogenic effect of IL-6 occurs selectively in a specific developmental window characterized by the presence of postmitotic neurons where synaptic contacts are not formed yet, thereby representing a vulnerable period in which IL-6 is specifically able to enhance the intrinsic capacity of neurons to form glutamatergic synapses. Indeed, when applied at later stages of neuronal development the cytokine, we found no effect at glutamatergic synaptic transmission and, accordingly, it has been reported that acute application of comparable IL-6 concentration on mature brain slices shifts the E/I balance through a reduction of GABAergic synaptic transmission without any alterations of glutamatergic inputs (Garcia-Oscos et al, 2012). These evidences highlight the capacity of IL-6 to affect distinct subpopulations of neurons at specific developmental stages, resulting in different effects on neuronal functioning.

The increased number of glutamatergic contacts, together with the lack of effect on inhibitory synapses is indicative of an E/I imbalance produced by IL-6. A proper E/I ratio is a fundamental factor ensuring a correct brain functioning in adulthood (Bhatia et al, 2019; Deco et al, 2014; Jelitai et al, 2016), and represents a pathological feature of different neurodevelopmental disorders, such as ASD and schizophrenia (Lisman, 2012; Nelson & Valakh, 2015; O’Donnell et al, 2017; Sohal & Rubenstein, 2019). To note, several mouse models of neurodevelopment disorders characterized by an E/I imbalance (Durand et al, 2007; Lee et al, 2015; Sala et al, 2001) also display altered brain functional connectivity (Ajram et al, 2017; Filipello et al, 2018; Pagani et al, 2019; Zhou et al, 2019), so highlighting the association between these two main hallmarks in brain disorders. Interestingly, we found an overall hyper-connectivity in the brain of adult mice prenatally exposed to IL-6, with a particular involvement of the hippocampal and somatomotor-thalamic regions. The relevance of this finding is underlined by recent results obtained in humans showing a tight association between maternal IL-6 elevations during pregnancy and altered brain connectivity in the newborns (Rudolph et al, 2018; Spann et al, 2018). Hence, our finding that IL-6 exerts a pro-synaptogenic role on glutamatergic synapses, associated with altered brain connectivity, may represent a possible molecular underpinning of the IL-6 effect detected in humans.

It has been previously found that IL-6 elevations in models of prenatal immune activation, engage other proinflammatory cytokines, such as IL-17, (Choi et al, 2016; Shin Yim et al, 2017) through the activation of maternal gut microbiota (Kim et al, 2017), which in turn leads to brain developmental defects (Choi et al, 2016; Shin Yim et al, 2017). Our evidence that the direct injection of IL-6 into the ventricles of the fetal brain at E15 increases the number of glutamatergic contacts in the adult offspring at P15, thus recapitulating the effects produced by maternal IL-6 elevations, indicates that IL-6 affects synapse formation directly in the embryos, without requiring the activation of other immune-related molecules in the mother. In line with this hypothesis, it is known that IL-6 can cross the placental barrier (Dahlgren et al, 2006), thus further supporting the notion that a maternal IL-6 elevation *per se* can directly reach the fetal brain thereby acting onto neurons.

Since the IL-6 receptor is expressed in the brain (Aniszewska et al, 2015; Gadient & Otten, 1993; Gadient & Otten, 1996; Rothaug et al, 2016) including in hippocampal neurons (Gadient & Otten, 1994; Sawada et al, 1993; Vereyken et al, 2007), a possible direct action of the IL-6 onto neurons might be expected. We provided robust evidences demonstrating that the effect of IL-6 on excitatory synapses was not ascribable to any indirect effect of the cytokine on brain cell types other than neurons. Indeed, IL-6 treatment enhanced excitatory transmission in virtually pure neuronal cultures, devoid of astrocytes. Furthermore, we ruled out any possible mitogenic effect of IL-6 on glial cells. This latter result is not trivial, given the crucial role played by IL-6 and STAT3 in different brain tumors (Lee et al, 2010; Levison et al, 2000; Selmaj et al, 1990; Weissenberger et al, 2004; Yu et al, 2009). However, the mitogenic action of the cytokine is driven by a chronic inflammatory state (Jones & Jenkins, 2018; Mantovani et al, 2008), whilst our paradigm relies on a transient activation of IL6-mediated signaling. Moreover, the cytokine did not promote any pro-survival effect on hippocampal neurons, as instead reported in other neuronal cell types, like catecholaminergic (Kushima et al, 1992); cholinergic (Hama et al, 1989) and retinal ganglion neurons (Mendonca Torres & de Araujo, 2001). Also, IL-6 did not affect neuronal excitability thus excluding indirect homeostatic compensatory effects such as synaptic scaling. All these results support the view of a direct action of IL-6 onto neurons resulting in a pro-synaptogenic effect at excitatory synapses. Furthermore, the effect appears to be highly specific for this immune molecule, as other key proinflammatory cytokines activated upon inflammation (e.g. INFγ, TNFα and IL1β), failed in promoting the enhancement of glutamatergic neurotransmission. IL-6 thus plays a unique role in the control of excitatory synaptogenesis and probably brain connectivity.

The long-lasting effect produced by IL-6 on glutamatergic synaptogenesis results from a transcriptional mechanism associated with a genomic rearrangement, and we identified the transcription factor STAT3 as a central molecular player underlying this effect. Although already studied in many different cellular context, including immune (Jiang et al, 2014; Maritano et al, 2004), cancer (Yu et al, 2009), neuronal progenitor (Gallagher et al, 2013) and in mature neurons (Fang et al, 2013; Leibinger et al, 2013a; Murase et al, 2012; Nicolas et al, 2012; Park et al, 2012), the direct role of STAT3 in neuronal synapse formation has never been investigated before. We show that STAT3 is both transcriptionally and functionally activated upon IL-6 elevation. To note, the IL6-dependent upregulation of the protein occurs transiently in a STAT3 independent manner, in contrast to immune cells in which STAT3 expression is autoregulated (Ichiba et al, 1998; Narimatsu et al, 2001). Importantly, STAT3 activation is causally linked with the increase of glutamatergic synapses and it promotes glutamatergic synaptogenesis in a cell autonomous fashion. We also demonstrate that the effect of STAT3 is mediated by transcriptional mechanisms (Heinrich et al, 1998), and does not rely on non-genomic effect as previously reported for synaptic plasticity (Nicolas et al, 2012).

It is worth noticing that STAT3 is significantly upregulated by IL-6 in two specific neuronal cell clusters, distinct from GABAergic neurons, thus indicating a predominant effect of the cytokine on *bona-fide* glutamatergic neurons, which likely explains the selective action of the cytokine at excitatory synapses.

One major accomplishment of our study is the identification of RGS4 as a STAT3 downstream neuronal gene, responsible for the boost of excitatory synapses through single cell transcriptomics. *RGS4* belongs to a class of gene family composed by 30 different members involved in the regulation of G-coupled receptor-associated signaling through their role as GTPase-activator proteins (GAP) in modulating intracellular second messengers (Bansal et al, 2007; Berman et al, 1996a; Berman et al, 1996b; Hepler et al, 1997). RGS4 is the most abundant isoform in the CNS, being highly expressed in prefrontal cortex, hippocampus thalamus and striatum (Ni et al, 1999; Nomoto et al, 1997). Accordingly, the single cell analysis revealed a high enrichment of RGS4 in neuronal specific cluster genes. Although RGS4 role has been well studied in mature neurons, whereby it regulates multiple aspects of neuronal physiology including synaptic transmission and plasticity (Gerber et al, 2016), only few studies have investigated the possible involvement of RGS4 in neuronal development (Cheng et al, 2013; Pallaki et al, 2017). Furthermore, its transcriptional regulation is completely unknown. We provide evidences that RGS4 expression is dependent on STAT3 genomic activity and its activation is crucial for the IL6-mediated boost of excitatory synapses. Given the importance of G-protein signaling in neuronal development (Munno et al, 2003; Shelly et al, 2010), we propose RGS4 as a new player in the process of glutamatergic synaptogenesis, although its precise role in this phenomenon needs further investigations. We hypotheses that an early upregulation of the protein might modulate G-protein dependent intracellular signaling thus affecting the intrinsic capacity of neurons to form excitatory synapses. The role of RGS4 in neuronal development is of particular interest in view of its involvement in neurodevelopmental disorders. Indeed, polymorphisms in RGS4 gene (Chowdari et al, 2002; Shirts & Nimgaonkar, 2004; Talkowski et al, 2006), and alterations of the protein levels (Dean et al, 2009; Erdely et al, 2006; Schwarz, 2018), were detected in patients affected by schizophrenia, thus making RGS4 one of the most promising candidate gene for this neurodevelopmental disorder (Levitt et al, 2006). It is now well established that autism and schizophrenia are two neurodevelopmental pathological conditions characterized by an aberrant synaptic connectivity (Frankle et al, 2003; Glausier & Lewis, 2013; Konopaske et al, 2014; Penzes et al, 2011). Given the importance of maternal Immune activation as a risk factor in neurodevelopmental disorders (Bauman et al, 2014; Estes & McAllister, 2016; Giovanoli et al, 2016; Lipina et al, 2013; Malkova et al, 2012; Missault et al, 2014), our findings that STAT3 and RGS4 expression are upregulated in the brain of embryos exposed to MIA represent a strong indication for a critical involvement of these two genes in inducing long-term consequences on synaptic density and brain connectivity upon prenatal inflammatory conditions.

Hence, our study provides a new piece of evidence about the complex cross-talk between immune and central nervous systems (Pozzi et al, 2018), unveiling the crucial contribution of two fundamental molecular neuronal players, STAT3 and RGS4, in abnormal synaptic formation and brain connectivity upon early IL6-mediated inflammation. Given the current lack of pharmacological tools for the prevention of neurodevelopment disorders, these results represent a promising perspective in the identification of new druggable pathways for early interventional strategies.

## Acknowledgements

We thank Prof. Valeria Poli, Department of Molecular Biotechnology and Health Sciences, University of Turin, Italy for kindly providing us all plasmids coding for the phosphomimetic mutants of STAT3; Dr Marinos Kallikourdis for Intellectual inputs and comments on the manuscript.

## Autor contribution

F.M. performed and analyzed electrophysiological, biochemical and immunofluorescence experiments; G.D. Performed and analyzed qPCR experiments; G.F. performed in vitro immunofluorescence experiments and analysis; M.R. and F.M. performed the in vivo injections on pregnant mice; S.M. and S.L performed the intraventricular injections and immunofluorescence analysis; C.P. performed single cell sequencing experiment; P.K. and A.T. analyzed and interpreted the single cell sequencing data; R.M. performed the ex vivo electrophysiological recordings; Ma.M., C.G. and V.Z. functional connectivity experiments and data analysis; E.M. acquisition and analysis of immunofluorescence experiments; M.M., D.P., E.M., F.M., G.D., designed, analyzed and interpreted the data. M.M. and D.P. conceived the study and wrote the manuscript.

## STAR Methods

### EXPERIMENTAL MODELS AND SUBJECT DETAILS

#### Animals, MIA model and *in utero* intraventricular injection

All experiments were performed using mice C57BL/6J (Charles River Laboratories) according the guidelines established by the European Community Council (Directive 2010/63/EU of September 22nd,2010) and were approved by the Institutional Animal Care and Use Committee (IACUC, permission number 467 and 565) of the Humanitas Research Hospital and by the Italian Ministry of Health. Mice were housed in a Specific Pathogen Free (SPF) facility under constant temperature (22 ± 1 °C) and humidity (50%) conditions with a 12 h light/dark cycle and were provided with food and water ad libitum. At gestational day 15, pregnant mothers were intraperitoneally (IP) injected with either 5 μg of IL-6, polyinosinic:polycytidylic acid (Poly(I:C)) 2 or 20mg/kg, or laparotomy was performed to inject 1 ul of IL-6 at 10ng/μl directly into the embryos lateral ventricles using the Nanoject II Auto-Nanoliter Injector (Drummond, cat.# 3-000-204). Saline solution was used as a control of injection. Only male embryos and offspring were selected for the ex-vivo and in-vivo analysis.

#### Primary cultures and cell lines

Primary hippocampal neurons were established from E18 C57BL/6 mice as previously described (Pozzi et al, 2013). Briefly, hippocampal regions were isolated from the brain in HBBS 1X (Hank’s Balanced Salt Solution) (Life technology), 1% Pen/Strep, 10mM Hepes) and, after trypsinization, they were dissociated and plated onto 24 mm-diameter round glass coverslips, previously coated with 0.1% Poly-L-Lysine (Sigma-Aldrich) in Borate buffer (50 mM Boric Acid, 15 mM Borax) pH 8.5. 80000 cells were seeded onto each coverslip. Cultures were grown in Neurobasal medium supplemented with 2% B27 and 1% Glutamax (Life Technology) at 37°C and 5% CO_2_. Neurons were transfected with Lipofectamine 2000 (Life Technology) at 5 days in vitro (DIV) according to the manufacturer’s protocol.

Neuro2a (N2A; ATCC^®^ CCL-131^™^) Neuroblastoma cell line was used to perform FACS experiments. N2A cells were cultured in complete DMEM medium (Life Technology) supplemented with 10% FBS, 1% pen/strep; 1% Ultraglutamine (Life technologies) until reaching the sub-confluence and then seeded in 60mm-diameter round dishes at 5×10^^5^ density.

### METHOD DETAILS

#### Drugs and Reagents

According to the type of experiment, the following reagents have been used: IL-6 (Peprotech), Poly(I:C) (Sigma-Aldrich), Galiellalactone (BioAustralis), CCG-63802 (Sigma-Aldrich), Stattic, Bicuculline, 6-Cyano-7-nitroquinoxaline-2,3-dione (CNQX), Tetrodotoxin (TTX), D-2-amino-5-phosphonovalerate (APV) (Tocris Bioscience).

#### Calcium imaging

Calcium imaging experiments were performed as previously described (Bedogni et al, 2016; Pozzi et al, 2013). Cultured neurons were loaded with the calcium sensitive dye Oregon Green 488 BAPTA 1-AM (Molecular Probes) for 1 h at 37°C in Neurobasal Medium and then imaged for calcium response. Electrical field stimulation was performed in KRH solution containing in mM: 125 NaCl; 5 KCl; 1,2 MgSO_4_; 1,2 KH_2_PO_4_; 25 HEPES; 6 Glucose; 2 CaCl_2_; pH 7,4.in the presence of CNQX 20 μM, APV 50 μM and Bicuculline 20 μM using a stimulation chamber (Warner Instruments, Hamden, CT). Electrical-evoked calcium transients were induced with a stimulus train of 40 stimuli (duration 1 msec; amplitude 90 mA) at 20Hz as previously reported (Pozzi 2013), using a train generation unit (Digitimer Ltd, DG2A) connected to a stimulus isolation unit (SIU-102; Warner Instruments, Hamden, CT). Recording chambers were placed on the stage of an IX-71 inverted microscope (Olympus, Hamburg, Germany) equipped with an EMCCD (electron-multiplying CCD) camera (Quantem 512×512, Photometrics). Illumination was obtained using a light-emitting diode LED (Cairn research, Optoled Lite), with a 20X objective. Regions of interest (ROIs) of about 15-pixel area were drawn on the cell cytoplasm of virtually all the cells in the recorded field. Time-lapse recording of calcium dynamics was performed with an acquisition rate of 5 Hz for 600seconds and off-line analyzed with MetaFluor software (Molecular Devices). Calcium responses were measure as ΔF (Fmax-F0) compared to the baseline (F0). All values were normalized to WT neurons at the same developmental stage within the same experiment. Cumulative data were then analyzed through Kolmogorov-Smirnov statistic to verify non-parametric distribution.

#### Electrophysiology

##### Ex vivo acute hippocampal slices

C57BL6 male mice at P15 were deeply anesthetized with isofluorane inhalation and decapitated. Brains were removed and placed in ice-cold solution containing the following (in millimolar): 87 NaCl, 21 NaHCO3, 1.25 NaH2PO4, 7 MgCl2, 0.5 CaCl2, 2.5 KCl, 25 D-glucose, and 7 sucrose, equilibrated with 95% O2 and 5% CO2 (pH 7.4). Coronal slices (300 μm thick) were cut with a VT1000S vibratome (Leica Microsystems) from medial Prefrontal Cortex (PFC). Slices were incubated at room temperature for at least 1 h, in the same solution as above, before being transferred to the recording chamber. During experiments, slices were superfused at 2.0 mL/min with artificial cerebrospinal fluid (ACSF) containing the following (in millimolar): 135 NaCl, 21 NaHCO3, 0.6 CaCl2, 3 KCl, 1.25 NaH2PO4, 1.8 MgSO4, and 10 D-glucose, aerated with 95% O2 and 5% CO2 (pH 7.4). Cells were examined with a BX51WI upright microscope (Olympus) equipped with a water immersion differential interference contrast (DIC) objective and an infrared (IR) camera (XM10r Olympus). Neurons were voltage (or current) clamped with a Multiclamp 700B patch-clamp amplifier (Molecular Devices, Union City, CA) at room temperature. Low-resistance micropipettes (2-3 MΩ) were pulled from borosilicate. The cell capacitance and series resistance were always compensated. Experiments in which series resistance did not remain below 10 MΩ (typically 5--8 MΩ) were discarded. Input resistance was generally close to 100-200 MΩ. Signals were low-pass filtered at 2 kHz, sampled at 10kHz and analyzed with Digidata 1440A (Molecular Devices). Recordings were made from cortical layer V pyramidal neurons. Excitatory and inhibitory synaptic basal transmission were recorded at - 70 mV and +10 mV respectively in the presence of 1 μM TTX, using the following pipette internal solution (in mM): 138 Cs-gluconate, 2 NaCl,10 HEPES, 4 EGTA, 0.3 Tris-GTP and 4 Mg-ATP (pH 7.2).

##### In vitro primary hippocampal neurons

Patch-clamp recordings were performed in an extracellular solution with the following composition (in mM): 130 NaCl, 5 KCl, 1.2 KH_2_PO_4_, 1.2 MgSO_4_, 2 CaCl_2_, 25 HEPES, and 6 Glucose, pH 7.4, with glass pipettes of 4-6 MΩ as recording electrodes. mEPSCs were recorded in the presence of bicucullin (20 μM), APV (50 μM) and tetrodotoxin (TTX; 1 mM) using the following internal solution (in mM): 135 K-gluconate, 5 KCl, MgCl_2_, 10 HEPES, 1 EGTA, 2 ATP, 0.5 GTP, pH 7.4. mIPSCs were recorded in the presence of 6-cyano-7-nitroquinoxaline-2,3-dione (CNQX; 20 μM), (2R)-amino-5-phosphonovaleric acid (APV; 50 μM) and TTX (1 mM) using the following internal solution (in mM): 68 K-gluconate, 68 KCl, 2 MgSO_4_, 20 HEPES, 2 ATP, 0.5 GTP, pH 7.2. Both mEPSCs and mIPSCs were recorded at - 70 mV as holding potential. Short term plasticity was evaluated through paired whole-cell recording as previously described (Liu et al, 2014; Maximov et al, 2007) using the following internal solution (in mM): 135 K-gluconate, 5 KCl, MgCl_2_, 10 HEPES, 1 EGTA, 2 ATP, 0.5 GTP, and 10 QX-314, pH 7.4. Presynaptic stimulation was achieved through a bipolar electrode (FHC, Concentric bipolar electrode Cat# CBAEC75) placed at 100-150 mm from the recorded neuron with 0,9 mA, 1 msec current injection through a stimulus isolation unit (SIU-102; Warner Instruments, Hamden, CT). Electrical signals were amplified by a Multiclamp 200 B (Axon instruments), filtered at 5 kHz, digitized at 20 kHz with a DIGIDATA 1440 and stored with pClamp 10 (Axon instruments). The resting potential was calculated at I=0 in current clamp configuration, whereas input resistance was calculated in voltage clamp configuration by using the slope of the I/V relationship of the steady state current measured at different hyperpolarizing voltage steps (from −100 to −70 mV). Only cells with an access resistance <20 MΩ were considered for the analysis of mEPSCs and mIPSCs, and <10 MΩ for the analysis of short-term plasticity. Access resistance was continuously monitored during the experiment and those cells in which access resistance was changed of more than 10% were rejected. The analysis of both mEPSC and mIPSC were performed with Mini analysis (Synaptosoft Inc., Fort Lee, NJ, USA) whereas the analysis of short-term plasticity was performed with Clampfit (Axon instruments).

#### Biochemistry

For total protein extraction, samples were homogenized using the following lysis buffer composed by 1% sodium dodecyl sulphate (SDS), 10mM Hepes at pH 7.4 and 2 mM EGTA and protein concentration was estimated using Bicinchoninic Acid Assay (BCA) kit (Thermo Fischer Scientific) as 20μg. Proteins were loaded with 2x Loading Buffer (100 mM Tris-HCl at pH 6.8; 4% SDS; 20% Glycerol; 200 mM 2-Mercaptoethanol, 2 mg Bromophenol-Blue) and fractionated by SDS-PAGE, then transferred to a nitrocellulose membrane using a transfer apparatus according to the manufacturer’s protocols (Bio-Rad). Membranes were stained with the following primary antibodies: mouse anti-STAT3 (Cell Signalling, 1:1000), rabbit anti-STAT3 P-Tyrosine 705 (Cell Signalling, 1:1000), mouse anti-STAT3 P-Serine 727 (Cell Signalling, 1:1000), rabbit anti-GAPDH (Synaptic System, 1:4000), mouse anti-PSD95 (UC Davis/NIH NeuroMab Facility, CA; 1:1000), guinea pig anti-vGLUT1 (Synaptic System, 1:1000), rabbit anti-Shank2/3 (Synaptic System, 1:1000), mouse anti-GAP43 (Millipore, 1:1000), rabbit anti-NFKB (Cell Signalling; 1:1000), rabbit anti-SNAP25 (Synaptic System, 1:1000), mouse antip65 (Cell Signalling; 1:1000). Immunodetection was performed with Clarity ™ Western ECL Substrate (Bio Rab) and analyzed through Chemidoc apparatus via ImageLab software (Bio-Rab).

#### Immunofluorescence analysis

##### Hippocampa/s/ices

Mice at P15 and P30 were deeply anesthetized with chloral hydrate (4%; 1 ml/100 g body weight, i.p.) and transcardially perfused with 4% paraformaldehyde (PFA). Post-fixed brains were collected and immunohistochemistry was performed on 50-μm hippocampal coronal sections with specific primary antibodies followed by incubation with the secondary antibodies. The following primary antibodies were used: guinea pig anti-vGLUT1 (1:1500; Synaptic Systems); mouse anti-V-gat (Synaptic Systems, 1:1000), rabbit anti-IBA (WAKO, 1:200), mouse anti-GFAP (Sigma-Aldrich, 1:400), guinea pig anti-NeuN (Synaptic System, 1:500), mouse anti-SATB2 (AbCam,1:100), rabbit anti-NeuroD2 (AbCam, 1:1500). Embryos’ brains (EB) were collected at GD16, 24 hours post mother injection, and post-fixed with PFA 4% for 16 hours. EB were embedded in 4% Low-Melting Agarose and 40μm-thick slices were cut. Brain slices underwent immunofluorescence staining against mouse anti-STAT3 (Cell Signalling, 1:1000). Secondary antibodies conjugated with Alexa-488, Alexa-555, or Alexa-633 fluorophores (Invitrogen) were used. All slices were counterstained with Hoechst-33342 (ThermoFisher) and mounted with Fluorsave (Calbiochem, San Diego, CA, USA).

##### In vitro primary hippocampal neurons

Neuronal cultures were fixed in 4 % PFA, 4 % sucrose, 20 mM NaOH and 5 mM MgCl_2_ in PBS, pH 7.4, for 8 minutes at room temperature (RT). Cultures were permeabilized and non-specific binding sites of proteins blocked with Goat Serum Dilution Buffer (GSDB; 15 % goat serum, 0.3 % Triton X-100, 450 mM NaCl, 20 mM phosphate buffer, pH 7.4) for 30 minutes. The following primary antibodies were used: guinea pig anti-vGLUT1 (Synaptic Systems, 1:1000), mouse anti-PSD95 (UC Davis/NIH NeuroMab Facility, CA, 1:800), mouse anti-ßeta III Tubulin (Promega Corporation, 1:800), rabbit anti-tubulin (Sigma-Aldrich, 1:80), rabbit anti-MAP2 (Millipore, 1:1000), mouse anti STAT3 (Cell Signalling, 1:1000). Secondary antibodies conjugated with Alexa-488, Alexa-555, or Alexa-633 fluorophores (Invitrogen) were used. Images were acquired using a Leica SP8I confocal microscope equipped with an ACS APO 40x oil immersion objective. Whole brain images were taken with DMi8 inverted light microscope equipped with ACS APO 10x dry objective (Leica Microsystems, Solms,Germany). Image analyses were performed using Bitplane Imaris 7.4 software (Bitplane AG, Zurich, Switzerland) and Fiji-ImageJ software (NIH,Bethesda, MD, USA).

#### Magnetic resonance imaging

N=16 mice (9 IL-6 and 7 vehicle control) underwent MRI to study the functional connectivity and white matter integrity of the entire brain. During the imaging sessions, the experimenters were blinded to the group. Data collection was performed on a Biospec 70/16 small animal MRI system (Bruker BioSpin) equipped with a cryogenic quadrature surface coil (Bruker BioSpin). For the detection of rs-fMRI we used a standard echo-echo gradient echo imaging sequence (GE-EPI, repetition time TR = 1 s, echo time TE = 15 ms, resolution in the plane RES = 0.22 × 0, 2 mm^2^, number of slices NS = 20, slice thickness ST = 0.4 mm, slice spacing SS = 0.1 mm, 900 volumes, sampling time = 15 min). In addition, diffusion-weighted images (DWI) were used to evaluate the structural integrity of the white matter (EPI spin echo multishot sequence, 4 segments, TR = 2 s, TE = 22 ms, RES = 0.2 × 0.2 mm^2^, NS = 28, ST = 0.4 mm, SS= 0 mm, values b = 0–1000 s / mm2, coding 94 directions, acquisition time = 9 min).

During MRI, the degree of anesthesia and the physiological parameters of the mouse were monitored according to a defined protocol to obtain a reliable measure of functional connectivity (Grandjean et al, 2014; Zerbi et al, 2015). Briefly, anesthesia was initiated with 4% isoflurane and the animals were intubated endotracheally and the tail vein was cannulated. The mice were placed on a compatible MRI base and artificially ventilated with 80 breaths / min, 1: 4 O_2_ / air ratio and 1.8 ml / h (CWE) flow. A bolus injection of medetomidine 0.04 mg / kg and pancuronium bromide 0.05 mg / kg was given and isoflurane was reduced to 1%. After 5 minutes, an infusion of medetomidine at 0.09 mg / kg / h and pancuronium bromide at 0.15 mg / kg / h were administered and the isoflurane was further reduced to 0.5%. The temperature of the animals was monitored using a rectal thermometer probe and kept in the cradle at 36.5 ° C ± 0.5 during measurements with a water heating system.

#### Resting-state fMRI

The datasets were preprocessed using an existing pipeline to eliminate unwanted time-series confusion, according to the guidelines of the Human Connectome project adapted to the mouse (Zerbi et al, 2015). In short, each rs-fMRI data set was entered into MELODIC (Multivariate Exploratory Linear Optimized Decomposition of Independent Components; (Beckmann & Smith, 2004)) to perform an independent component analysis (ICA, number of components set to 60). This included the correction and regression of head movement and in-plane smoothing with a 0.3 x 0.3 mm kernel. We used FSL-FIX for the removal of the variance of artifact components. The artifact-corrected data sets were then band-pass filtered (0.01-0.25 Hz), normalized in a GE-EPI study-specific template, and then to the Allen Common Coordinate Framework Version (CCF, v3) with Advanced Normalization Tools (ANTs v2.1, http://picsl.upenn.edu/software/ants/).

#### Diffusion MRI

For diffusion MRI, the preprocessing steps consisted of individual realignment of the diffusion images, followed by eddy current correction and tensor estimation as in (Zerbi et al, 2013a). From the eigenvalues of the diffusion tensor, fractional anisotropy (FA), mean diffusivity (MD), and first eigenvalue (λ1) maps were calculated. The resulting volumes were spatially normalized to the Allen CCF template using linear affine and nonlinear elastic transformations in ANTs and thereafter FA, MD, and λ1 values were extracted from seven major white matter structures: anterior commissure, fimbria, corpus callosum, fornix, cingulum, internal capsule, and cerebral peduncle.

#### Behavioural studies

Animals were kept under a light-dark cycle (12 light-12 dark) in a temperature and humidity controlled room. Room lights were low during all procedures. A camera was mounted above the arena and object exploration was tracked using a computer running the Smart Software (Panlab, Harvard Apparatus). Only males were used in these studies.

#### Open Field Analysis

For the open field, each subject was gently placed in the centre of the open field and allowed to explore freely all the arena for 10 min. At the end, the animal was removed and the arena cleaned with 70% ethanol and dried before testing the next animal. Locomotor activity was indexed as the distance spent in centre, periphery and total distance travelled.

#### Elevated plus maze

The Elevated Plus Maze (EPM) was conducted as previously described (Hagenbuch et al, 2006). Mice were allowed to freely explore the entire apparatus for 5 min. The percentage of time spent in the arms was scored as measure of anxiety-related behavior. The arm entry was defined as having all four paws into the arm of the EPM.

#### Object location Recognition and Novel Object Recognition test

The test was performed in the open-field apparatus. This test was used to assess short-time memory retention. Between trials, the objects and the box were cleaned with 70% ethanol. Mice were first habitued to an open field chamber by allowing free exploration of an empty chamber for 5 min. The Oject location and Novel object recognition test included three sessions. In the first session, two identical objects were placed in the testing arena and each mouse was allowed to explore the objects for 10 minutes to facilitate the familiarization. These objects presented similar textures, colors and sizes. One hour later, one of the identical objects was displaced and the mouse was placed back into the chamber and allowed to explore objects for 10 min (Object location Recognition phase). Lastly,1 hour later, the mouse was returned for 10 min in the chamber where a novel object replaced one of the 2 identical familiar objects (Novel Object Recognition Phase). The amount of time the mouse spent physically investigating each of the objects was manually determined for all the trials and was used to calculate the discrimination index.

#### Nissl staining

Brains were perfused with 4% PFA and 50 μm thick coronal slices were obtained at the vibratome. Slices were mounted on charged coverslips and let dry overnight. They were then re-hydrated with an alcohol scale from 100% to water and subsequently stained for 5 minutes in a 0.1% cresyl violet solution. Afterwards, slices were differentiated again in alcohol 95% for 5 minutes according to the intensity of the staining, de-hydrated in alcohol 100% and a subsequently fixed with a xylene-based medium. Images were taken at BX61VS Olympus microscope using a 10x/0,40 UPlanSApo objective.

#### Luciferase assay

To monitor the transcriptional activity of STAT3 cultured neurons were transduced using Cignal Lenti STAT3 reporter Kit (Qiagen) at 3 DIV and exposed to IL-6 and Galiellalactone for 48 hours at 10 DIV. Neurons were then processed according to manufacturer’s instructions and Firefly luciferase activity was detected with Luciferase assay system (Promega) and Synergy H2 (Biotek).

#### Quantitative RT-PCR

Cultured neurons were homogenized prior to RNA extraction in 500 μl of TRI-reagent (Zymo research). Total RNA was isolated using the RNA Direct-Zol™ MiniPrep Isolation Kit (Zymo research) according to the manufacturer’s guidelines. The RNA was eluted in 25 μL DNase/RNAse-free water, quantified using NANOdrop 2000c spectrophotometer (Thermo Fisher Scientific) for RNA concentration and 260/280 nm optical density ratios. Reverse transcription was performed using 1 μg RNA with a High Capacity cDNA RT kit (Applied Biosystems). Quantitative real-time polymerase chain reaction (qRT-PCR) was performed with Sybr Green detection kit (SensiFAST SYBR Lo-ROX, Bioline) with RT-PCR Viia7 software system (Applied Biosystems) in a final volume of 10 μl. Each gene was subjected to at least duplicate measurements and data analyses were performed with the comparative ΔΔC_T_ method. The RNA levels were normalized against *gapdh* and *actin3* as indicated. The following Sybr oligos were used*: gapdh* Fw: TTCCAGAGGGGCCATCCACAG, Rv: GGTCACCAGGGCTGCCATTTG; *actin* Fw: GCCATCCTGCGTTCTGGA, Rv: GCTCTTCTCCAGGGAGGA*; Rgs4* Fw: GTCGGAATACAGCGAGGAGAAC, Rv: GGAAGGATTGGTCAGGTCAAGATAG; *Stat3* Fw: CCATGCTGAGCATCGAGCAGCTGACA; Rv: TCA CACAGATGAACTTGGTCTTCAGG.

#### Fluorescence Activated Cell Sorting (FACS) analysis

The day after plating N2A were co-transfected according to three experimental conditions with the following plasmids: 1) pCDNA 3.1 EGFP; 2) pCDNA 3.1 EGFP + pCDNA 3.1 myc 768-STAT3 Y795E; 3) pCDNA 3.1 EGFP + pCDNA 3.1 myc 768-STAT3 Y705F and maintained for 6 hours in the transfection medium. After 24 hours, they were gently detached with Accutase solution (Sigma-Aldrich) for 3 minutes at 37°C, centrifuged at 500 rcf for 10 minutes and then sorted out by means of BD FACS Melody™ cell sorter (BD, Biosciences). GFP positive (+) and GFP negative (-) cells were collected for each condition, lysed with TRI-Reagent and subsequently processed for total RNA extraction and qRT-PCR.

#### Single cell sequencing

Hippocampal neurons untreated and treated with IL-6 (10ng/ml at 1 DIV and 4 DIV) were collected at 5 DIV using Accutase solution (Sigma-Aldrich) and resuspended in 1ml of PBS -/- with 0.04% BSA (Sigma Aldrich), centrifuged at 450 rcf for 7min and washed twice with PBS-/- with 0.04% BSA. After the second wash, cells were resuspended in 30 ul and counted with an automatic cell counter to get a precise estimation of total number of cells recovered. ~5,000 cells were loaded into one channel of the Single Cell Chip A for each sample using the Single Cell 3’ v2 single cell reagent kit (10X Genomics) for Gel bead Emulsion generation into the Chromium system. Following capture and lysis, cDNA was synthesized and amplified for 14 cycles following the manufacturer’s protocol (10X Genomics). 50 ng of the amplified cDNA were then used for each sample to construct Illumina sequencing libraries. Sequencing was performed on the NextSeq500 Illumina sequencing platform following 10x Genomics instruction for reads generation.

#### Single cell mapping and clustering

Raw sequencing data (bcl-files) were converted to fastq files with Illumina bcl2fastq tool, integrated into the CellRanger (10X Genomics) suite (version 2.1.1). The CellRanger analysis pipeline was used to generate a digital gene expression matrix staring from raw data. Pre-build mouse genome (version mm10-1.2.0) was used as genome reference. CellRanger count module was used to map reads with default settings setting and sequence length set to r1-length=26 and --r2-length=50. At least 90,000 reads per cell were produced. The raw digital gene expression matrix (UMI counts per gene per cell) was imported in R https://www.R-project.org/ version 3.5.2 using Seurat R package (version Seurat_2.3.4) (Butler et al, 2018). Briefly, UMI counts per gene per cell for each biological replicate was imported in Seurat. Initial quality control in each biological replicate was assessed, by filtering out cells meeting any of the following criteria: less than 200 or more than 10,000 unique genes expressed, more than 50,000 UMIs, or more than 15.0% of reads mapping to mitochondria. Data was normalized through a global-scaling method, converted by a scale factor (10,000 by default) and log-transformed. Scaling the data and removing unwanted sources of variation was applied through ScaleData Seurat function. Detection of variable genes across the single cells in each sample was performed. The resulting gene list was used to perform a canonical correlation analysis (CCA) using the first 30 canonical vectors. After aligning subspaces, clustering was performed through FindClusters function, using the first 20 dimensions. Uniform Manifold Approximation and Projection (UMAP) dimensional reduction was computed with RunUMAP function.

Clusters enrichment was determined by running the AUCell Bioconductor R Package v1.2.4 (Aibar et al, 2017) with default parameters.

AUCell evaluates each cell individually. Genes are ranked from highest to lowest expression value. The top 5% of the genes in the ranking are used to calculate the “Area Under the Curve” (AUC) value for the gene-set (signature) of interest. The AUC estimates the proportion of genes in the signature that are highly expressed within the cell. The distribution of AUC values across all the cells allows exploring the relative expression of the signature: cells expressing more genes from the gene-set will have values higher than cells expressing fewer.

To determine clusters of cells that are more likely of the cell type according to the gene signature, cells are split into two groups: cells that pass the assignment threshold (cells where the gene-set is defined as “active”), and cells that don’t (cells where the gene-set is defined as “non-active”). The assignment threshold was calculated automatically by AUCell by examining the AUC distribution.

Clusters enrichment was determined by running the AUCell Bioconductor R Package v1.2.4 (Aibar et al, 2017) with default parameters. Three gene-sets were considered: two included in the AUCell package: neurons and astrocytes from (Cahoy et al, 2008; Cembrowski et al, 2016; Lein et al, 2007), one manually-curated GABAergic gene signature from (Harris et al, 2018) and proliferating cells from (Kowalczyk et al, 2015; Tirosh et al, 2016).

A Complete list of gene signatures used for identifying each cluster was added in Table 1. Cells that pass the assignment threshold are colored in shades of pink-red while cells that don’t are colored in black-blue in Supplementary Fig. 8.

#### Promoter sequence analysis

The bioinformatic prediction for transcription factor binding sites in RGS4 promoter region was performed using JASPAR database (http://jaspar.genereg.net/) (Khan et al, 2018; Sandelin et al, 2004)

#### Single cell markers identification and differential expression

We identified markers specific to each cluster using FindAllMarkers Seurat function with the following settings (only.pos=TRUE, min.pct=0.05, thresh.use=0.05). We defined a gene to be a cluster-specific marker if it was differentially up-regulated in a specific cluster (Wilcoxon test), as compared to each of the remaining clusters. To identify differentially expressed genes within the same cluster in different experimental conditions and between different clusters in the same experimental condition, we used MAST R package (Finak et al, Genome Biology, 2015) implemented in Seurat FindMarkers function with an FDR cut-off of 0.05 and a logfc.threshold=0.20.

### QUANTIFICATION AND STATISTICAL ANALYSIS

The numerical data shown in the figures are presented as means ± SEM. The normal distribution of experimental data was assessed using Kolmogorov Smirnov and Shapiro-Wilk normality tests. If not specifically indicated, to compare two normally distributed sample groups, the Student’s unpaired two-tailed t-test was used. To compare two sample groups that were not normally distributed, we used Mann–Whitney’s non-parametric test. To compare more than two normally distributed sample groups, we used one ANOVA, followed by Tukey’s multiple comparisons test. To compare more than two groups that were not normally distributed, we used one One-way ANOVA on ranks followed by Dunn’s multiple comparison test. One sample t test or Wilcoxon Signed Rank Test were used in some experiments to analyze normally distributed or not normally distributed data set, respectively. Statistical analysis was performed by using SigmaPlot (Systat), GraphPad (Prism) or OriginPro (OriginLab) software.

### DATA AND CODE AVAILABILITY

The scRNA-seq data, together with global mapping statistics, has been deposited in GEO under accession code GEO: GSE130803

