## Supplemental Figures for "Transient Maternal IL-6 boosts glutamatergic synapses and disrupts hippocampal connectivity in the offspring"

**SUPPLEMENTARY FIGURE LEGENDS**

**Figure S1 related to Figure 2: Transient maternal elevation of IL-6 disrupts normal brain network connectivity within specific large-scale circuits.**

**(A)** Representative scheme showing the experimental procedure used for the *in vivo* approach: a single pulse of either vehicle (saline, as control) or IL-6 (5µg) was intraperitoneally injected in pregnant mothers at gestational day 15 and the offspring were analyzed at different post-natal day with different experimental techniques.

**(B)** Immunofluorescence analysis of Vglut-1 and V-gat positive puncta in CA1 hippocampal neurons at P30 in the two conditions. Scale Bar 10µm.

(**C)** Quantitative analysis of Vglut-1 and V-gat area (vehicle n=3 mice; IL-6 n=3 mice.). **p=0,0046.

**(D)** Circos-plot showing the anatomical location of hypo-connected edges (n=36) in IL-6 mice compared to vehicle-treated mice ($p$<0.05, uncorrected).

**(E)** Dual-regression analysis in 15 resting-state networks (RSNs) revealed a significant increase only in the Dorsal Hippocampal network strength in the IL-6 group compared to vehicle mice. Multivariate ANOVA, Bonferroni corrected across 15 RSNs.

**(F)** Fractional anisotropy (FA) was assessed by diffusion tensor imaging and quantified in nine large white matter structures. None of these structures show significant microstructural differences between IL-6 and vehicle-treated mice. Multivariate ANOVA, Bonferroni corrected across 9 structures.

**Figure S2 related to Figure 2: Transient maternal elevation of IL-6 does not alter the gross anatomy and the glial composition of the brain in offspring**

**(A).** Nissl staining of three independent P15 brain coronal sections, at different rostro-caudal levels, derived from mice prenatally exposed to Vehicle- or IL-6 via maternal intraperitoneal injection. Scale bar 500 µm. Ctx: Cortex, Str: Striatum, Hp: Hippocampus. LV: Later Ventricle. No gross anatomical malformations are present.

**(B).** Representative image of P30 coronal brain sections stained for Satb2 (green), NeuroD2 (red), NeuN (grey) Hoechst (blue), obtained from mice prenatally exposed to either Vehicle or IL-6. Scale bar 1000 µm. Relative image of the selected (dashed rectangle) area of the somatosensory cortex showed below. Scale bar 500 µm. Right panel: quantitative analysis of SATB2 intensity (Vehicle n= 4 mice, IL-6 n= 4 mice. Two independent experiments). Mann-Whitney test

**(C)** Immunofluorescent analysis of coronal brain section at P15 in CA1 hippocampal region, obtained from mice prenatally exposed to vehicle- or IL-6 via maternal intraperitoneal injection, stained with Hoechst (blue), GFAP (red), IBA1 (green), scale bar 50 µm.

**(D)** Quantitative analysis of astrocyte properties (left panels) including GFAP mean intensity and astrocyte density, and microglial parameters (right panel) including cell density, number of branching and junctions per cell, and length of total branches. (Vehicle n=3 mice, IL-6 n=3 mice.) Mann-Whitney test.

**Figure S3 related to Figure 3: Passive properties of neurons exposed to chronic treatment of IL-6 at 10ng/ml**

(A) Quantitative analysis of resting potential measured in current clamp configuration (I=0) in untreated and chronically treated neurons with IL-6 10 ng/ml. (B) Quantitative analysis of membrane resistance measured in voltage clamp configuration in untreated and chronically treated neurons with IL-6 10 ng/ml (Ctrl n=26 cell, IL-6 n=23 cells. Three independent experiments). Mann-Whitney test.

**Figure S4 related to Figure 3: IL-6 doesn't affect electrically evoked neuronal calcium transients and it acts in an astrocyte dependent manner without altering cellular composition of the cultures.**

(A) Immunostaining of neuronal cultures treated (lower panels) or not treated (upper panels) with cytosine arabinoside (Ara-C, 3 μM) using antibodies against MAP-2 (neurons, in red) GFAP (astrocytes, in green) and both (Hoechst, in blue). Ara-C treated neurons are virtually devoid of glial component. Scale Bar 100μm.

(B) Representative electrophysiological traces of mEPSC in Ara-C treated cultures both in control condition and upon the chronic treatment with IL-6 10 ng/ml.

(C) The quantitative analysis of amplitude and frequency of mEPSC in Ara-C treated neurons in both control and IL-6 treated neurons. (Ctrl n= 24, IL-6 n= 30. Three independent experiments) Mann-Whitney test. ***p= 0.0002; *p=0.0452;

(D) Representative electrophysiological traces of excitatory post-synaptic currents (EPSC) evoked by paired pulse at 50 msec both in control condition and upon IL-6 chronic treatment at 10 ng/ml.

(E) Short term plasticity measured as paired pulse ratio between first and second EPSC evoked at different interpulse intervals. (Ctrl n=15 cells, IL-6 15 cells. Three independent experiments). ANOVA followed by Tukey's multiple comparison test.

(F) Quantitative analysis of a single electrically evoked EPSC recorded in the two conditions. (Ctrl n=42 cells, IL-6 n= 42 cells. Three independent experiments). Mann-Whitney test. *p<0,0138.

(G) Pseudocolor images of cultured neurons loaded with calcium sensitive dye Oregon-Green and imaged (505 nm) in resting state (left panel) and upon an electrical field stimulation (90 mA for 2 sec @ 20Hz, right panel) in the present of synaptic transmission blockers (scale bar 50 μm).

(H) Temporal analysis of intracellular calcium changes during electrical stimulation measured at somato-dendritic level with time lapse imaging at a rate of 2 Hz.

(I) Quantitative analyses of intracellular calcium influx in control condition and upon chronic IL-6 10 ng/ml treatment. (Ctrl n=184 cells, IL-6 212 cells. Three independent experiments). Mann-Whitney test.

(J) Immunofluorescence analysis of cultured neurons untreated and chronically treated with IL-6 10ng/ml, stained with antibodies against specific neuronal (NeuN, green) glial (GFAP, red) and a general cellular marker (Hoechst, blue). Scale Bar 100μm.

(K) Quantitative analysis of the total number of cells identified through Hoechst staining. (Ctrl n= 32 coverslips, IL-6 n= 31 coverslips. Roughly 3-4 field analyzed for each coverslip. Three independent experiments) Mann-Whitney test.

(L) Evaluation of ratio analysis between GFAP positive (astrocytes) and NeuN positive (neurons) cells in the two conditions (Three independent experiments) Mann-Whitney test.

**Figure S5 related to Figure 6: Possible role of other proinflammatory cytokines as prosynaptogenic molecules. Effect of Stattic dosage on neuronal death and glutamatergic synaptic transmission in developing cultures.**

(A) Representative electrophysiological traces of mEPSCs recorded in cultured neurons at 14 Days in vitro (DIV) in control condition and upon a single application of distinct proinflammatory cytokines at 1 DIV.

(B) Quantitively analysis of mEPSCs frequency and amplitude in the indicated conditions. (ctrl n= 16 cells, INF$\gamma$ n=14 cells, TNF$\alpha$ n=10, IL1$\beta$ n=12 cells. Three independent experiments.) One-way ANOVA on ranks followed by Dunn's multiple comparison test.

(C) Neuronal death assay performed in neuronal cultures incubated at 1 DIV with different concentrations of Stattic (0; 0,5; 1; 2; 4 μM) using Calcein (live cells, green), Propidium Iodide (death cells, red) and Hoechst (total cells, in blue) in vivo staining. Scale bar 100 μm.(Lower panel) The percentage of dead (PI positive cells) and (Right panel) live cells (calcein positive cells) were evaluated as percentage of the total number of cells (Hoechst positive cells) (n= 9-15 coverslips analyzed for each condition. Roughly 100 cells analyzed for each coverslip. Three independent experiments.). One-way ANOVA on ranks followed by Dunn's multiple comparison test. PI: **p=0,003; ****p<0,0001. Calcein: *p=0,0149, ****p<0,0001.

(D) Representative traces of mEPSCs recorded in cultured neurons at 14 DIV in control condition and upon a single incubation of Stattic 1μM at 1 DIV. (Lower panel) Quantitative analysis of mEPSCs frequency and amplitude in the two conditions. (Ctrl n= 22 cells, IL-6 n=18 cells. Three independent experiments.) Mann Whitney test.

**Figure S6 related to Figure 5: The increase of STAT3 expression level is not due to STAT3 phosphorylation itself and occurs transiently.**

**(A)** Schematic representation of the experimental procedure: IL-6 was chronically treated throughout the *in* vitro development of hippocampal neurons, from 1 DIV up to 13 DIV, by adding the cytokine with either vehicle or Stattic 1 μM every 3 days (see Figure 3D).

**(B)** Representative traces of electrophysiological recordings of mEPSCs performed in neuronal cultures at 14 DIV in the indicated conditions.

**(C)** Quantitative analysis of mEPSCs frequency and amplitude in the indicated conditions. (Ctrl n=22 cells, IL-6 n=26 cells, IL-6 stattic n=31 cells. Four Independent experiments). One-way ANOVA on ranks followed by Dunn's multiple comparison test. Ctrl-IL6 ***p=0,0004; IL6-IL6 stattic***p=0,0002.

**(D)** Upper panel: western blot analysis of STAT3 expression level in cultured neurons at 14 DIV chronically treated with IL-6 together with either vehicle or Stattic 1 μM every 2 days (refer to scheme in A). Lower panel: quantification of the optical density of total STAT3 protein levels in the different conditions normalized by GAPDH level (Three independent experiments). One sample t test p*<0,05.

**(E)** Western blot analysis of a panel of synaptic and non-synaptic proteins (left panel) evaluated in cultured neurons at 14 DIV in control condition and upon chronic treatment of IL-6 with the relative quantitative analysis (right) normalized by GAPDH level. (Four independent experiment) One simple t test. *=p<0,05

**(F)** Schematic representation of the experimental procedure: IL-6 10ng/ml was incubated for a short period to cultured neurons, at 1 and 4 DIV (similar to scheme in Figure 2A) and collected at 14 DIV.

**(G)** Western blot analysis of STAT3 expression level (left panel) in cultured neurons incubated for a short period with IL-6 10ng/ml and analyzed at 14 DIV, (right panel) with the relative quantitative analysis normalized by GAPDH level. (Three independent experiments) Mann Whitney test.

**Figure S7 related to Figure 6 and 8: Dose-related effect of Galiellalactone and CCG-063802 on neuronal death performed in developing neurons.**

**(A)** Neuronal death assay performed in cultured neurons incubated with different concentrations of Galiellalactone (0; 2; 4; 8; 16 µM) at 1 DIV using Calcein (green), Propidium Iodide (red) and Hoechst (blue) in vivo staining. Scale bar 100 µm

**(B)**. The percentage of dead (PI positive cells, left panel) and live cells (calcein positive cells, right panel) were evaluated as percentage of the total number of cells (Hoechst positive cells*)* (n= 10-12 coverslips analyzed for each condition. Roughly 100 cells analyzed for each coverslip. Three independent experiments.). One-way ANOVA on ranks followed by Dunn's multiple comparison test. PI: ***p=0,008; Calcein: ***p=0,008.

**(C).** Western blot analysis STAT3 Phosphorylation at Tyrosine 705 in control condition and upon acute stimulation (30 min) of IL-6 in the presence of either vehicle or Galiellalactone at 1 and 4 µM.

**(D)** Representative traces (left panel) of mEPSCs recorded in cultured neurons at 14 DIV treated with Galiellalactone 4 µM, at 1 DIV, and the relative quantitative analysis (Ctrl n= 13 cells, Gall4 n= 15 cells. three independent experiments) Mann Whitney test.

**(E)** Neuronal death assay performed in neuronal cultures incubated with different concentrations of CCG-063802 (0; 1,25; 2,5; 5; 7,5; 15 mM) a 1 DIV using Calcein (green), Propidium Iodide (red) and Hoechst (blue) in vivo staining. Scale bar 100 µm

**(F)** The percentage of dead (PI positive cells, left panel) and live cells (calcein positive cells, right panel) were evaluated as percentage of the total number of cells (Hoechst positive cells*)* (n= 9-18 coverslips analyzed for each condition. Roughly 100 cells analyzed for each coverslip. Three independent experiments.). One-way ANOVA on ranks followed by Dunn's multiple comparison test. PI: **p=0,0145, ***p=0,0002, ****p<0,0001; Calcein: **p=0,0048, ****p<0,0001

**Figure S8 related to Figure 7: Cluster identification methodology and predictive analysis of *Rgs4* promoter region.**

**(A)** Results from clustering using Seurat R package with resolution parameters ranging from zero to 3. Red arrow indicates resolution used in this study.

**(B)** Left: UMAP projection of all cells coloured by AUC (Area under the curve) values of Neuron signature (Cahoy et al, 2008; Cembrowski et al, 2016; Lein et al, 2007). Pink-red: cells with active gene-set. Black-blue: cells with non-active gene-set. Right: Barplot of percentage of cells with active gene-set for each cluster.

**(C)** Left: UMAP projection of all cells coloured by AUC (Area under the curve) values of Astrocyte signature (Cahoy et al, 2008; Cembrowski et al, 2016; Lein et al, 2007). Pink-red: cells with active gene-set. Black-blue: cells with non-active gene-set. Right: Barplot of percentage of cells with active gene-set for each cluster

**(D)** Left: UMAP projection of all cells coloured by AUC (Area under the curve) values of GABAergic signature (Harris et al, 2018). Pink-red: cells with active gene-set. Black-blue: cells with non-active gene-set. Right: Barplot of percentage of cells with active gene-set for each cluster.

**(E)** Heat map showing the average expression value of G2/M and S cell cycle related genes signatures across all cells belonging to cluster 0 to 7. Average expression scale is shown on the right.

**(F)** Predicted Stat3 response element in the promoter of Rgs4, together with score and position. JASPAR[2018] scan tool was used to assess enrichment. Only hits with a relative profile score threshold above 80% was considered significant.

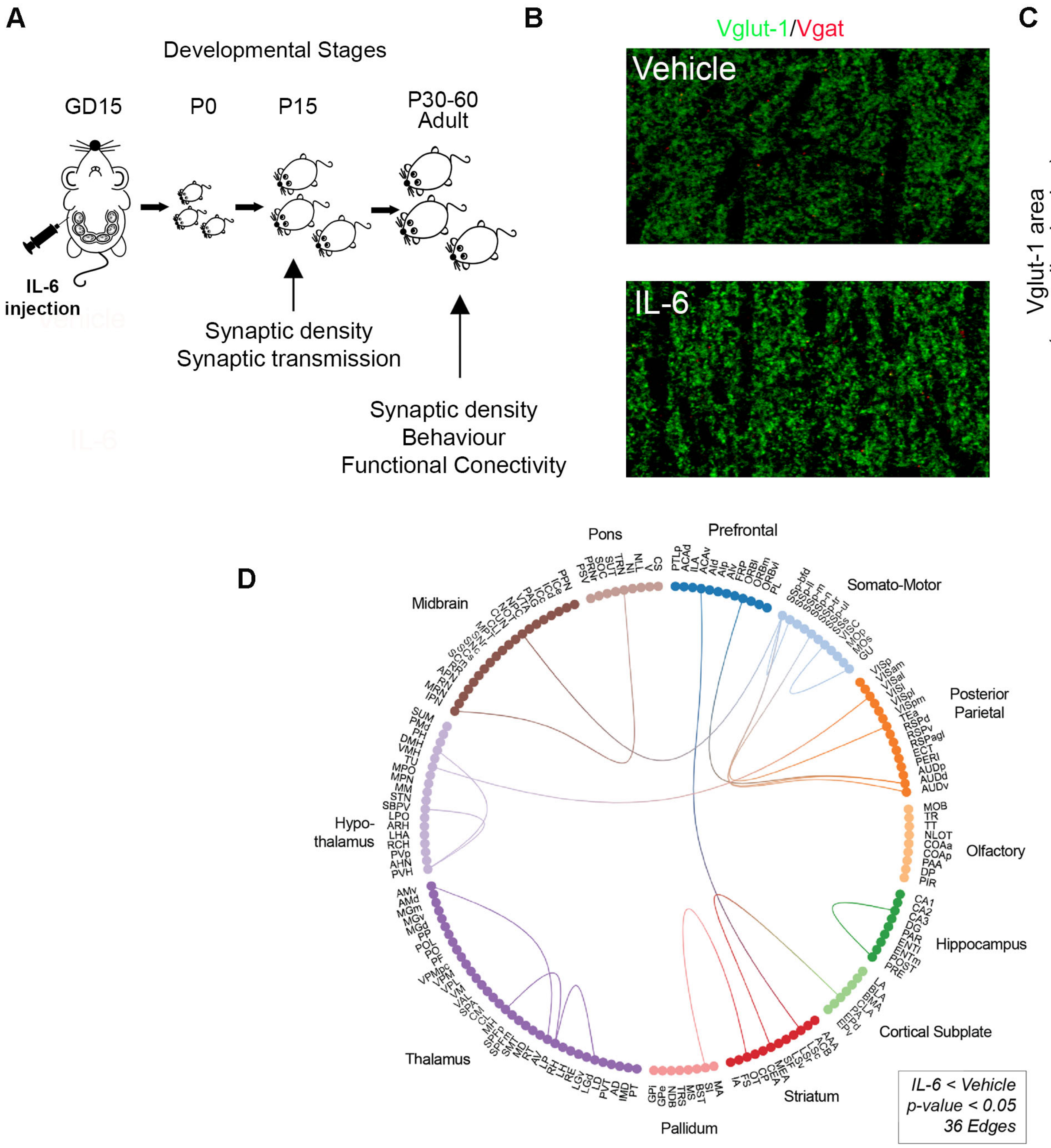


**E**

Pairwise Comparisons

| Independent Components | (I) Group | (J) Group | Mean Difference (I-J) | Std. Error | Sig.b | Network |
|---|---|---|---|---|---|---|
| IC_01 | CTRL | IL6 | 24.186 | 14.858 | 0.13 | Secondary Sensory (BFD) |
| IC_02 | CTRL | IL6 | -11.05 | 12.44 | 0.392 | Retrosplenial (DMN-like) |
| IC_03 | CTRL | IL6 | 13.454 | 10.045 | 0.205 | Lateral Striatum |
| **IC_04** | **CTRL** | **IL6** | **-8.239*** | **2.973** | **0.017** | **Dorsal Hippocampus** |
| IC_05 | CTRL | IL6 | 6.257 | 8.085 | 0.454 | Motor |
| IC_06 | CTRL | IL6 | 5.45 | 4.921 | 0.29 | Visual |
| IC_07 | CTRL | IL6 | 17.751 | 8.769 | 0.066 | Primary Sensory |
| IC_08 | CTRL | IL6 | 3.432 | 5.104 | 0.514 | Anterior Cingulate (Salience-like) |
| IC_09 | CTRL | IL6 | -9.495 | 5.147 | 0.09 | Auditory / Temporal Association |
| IC_10 | CTRL | IL6 | -0.098 | 1.117 | 0.931 | Amygdala |
| IC_11 | CTRL | IL6 | 1.453 | 1.307 | 0.288 | Olfactory / Piriform |
| IC_12 | CTRL | IL6 | 1.02 | 2.092 | 0.635 | Striato - motor |
| IC_13 | CTRL | IL6 | 0.528 | 0.731 | 0.484 | Ventral Hippocampus / Subiculum |
| IC_14 | CTRL | IL6 | 1.845 | 1.203 | 0.151 | Ventral Striatum |
| IC_15 | CTRL | IL6 | -0.454 | 0.646 | 0.496 | Thalamus |

Based on estimated marginal means
* The mean difference is significant at p-value<0.05
b Adjustment for multiple comparisons: Bonferroni.

**F**

| White Matter tract | (I) Group | (J) Group | Mean Difference (I-J) | Std. Error | Sig.b |
|---|---|---|---|---|---|
| Anterior Commisure | CTRL | IL6 | 0.014 | 0.033 | 0.672 |
| Fimbria | CTRL | IL6 | 0 | 0.008 | 0.981 |
| Corpus Callosum | CTRL | IL6 | 0.005 | 0.007 | 0.462 |
| Fornix | CTRL | IL6 | 0 | 0.01 | 0.973 |
| Cingulum | CTRL | IL6 | 0.006 | 0.01 | 0.57 |
| Ventral Hippocampal Commissure | CTRL | IL6 | 0.026 | 0.019 | 0.186 |
| Internal Capsule | CTRL | IL6 | -0.003 | 0.005 | 0.61 |
| Posterior Commissure | CTRL | IL6 | 0.011 | 0.01 | 0.3 |
| Cerebral Peduncle | CTRL | IL6 | 0.006 | 0.008 | 0.406 |

Based on estimated marginal means
* The mean difference is significant at p-value<0.05
b Adjustment for multiple comparisons: Bonferroni

Mirabella et al., Figure S1

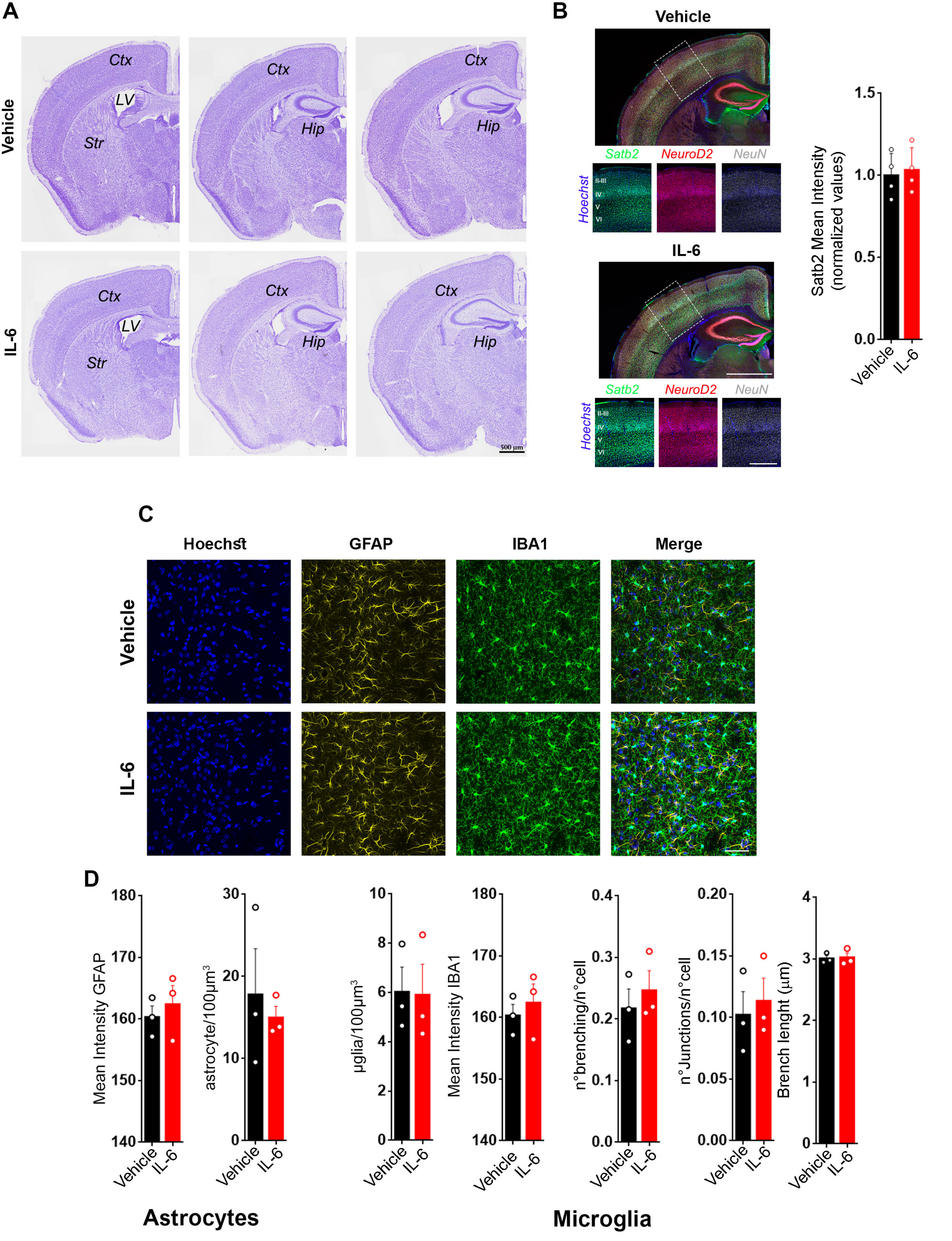


Mirabella et al., Figure S2

Mirabella et al., Figure S3

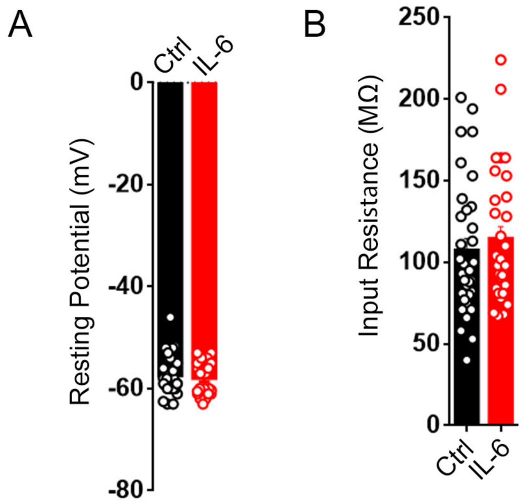

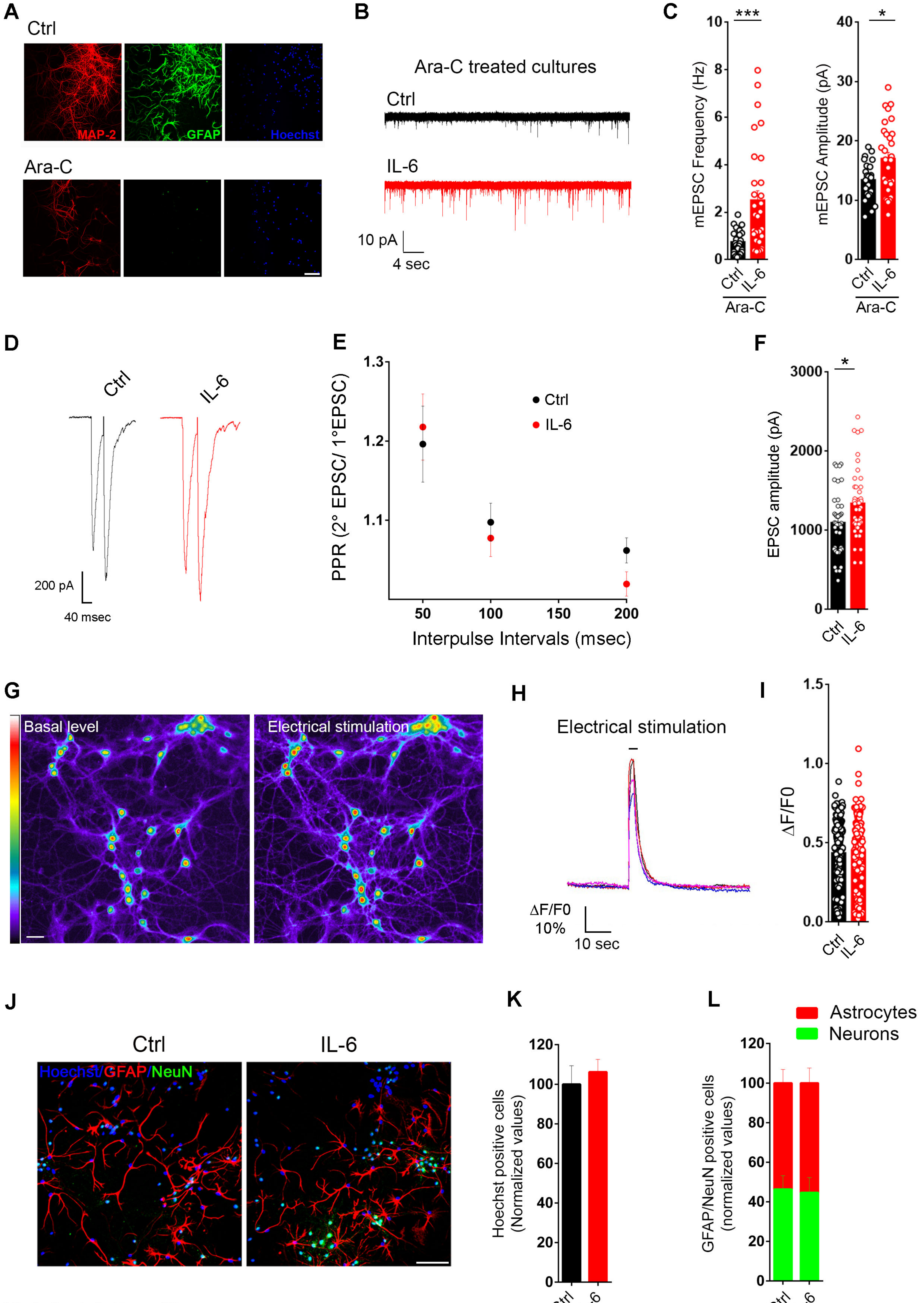


Mirabella et al., Figure S4

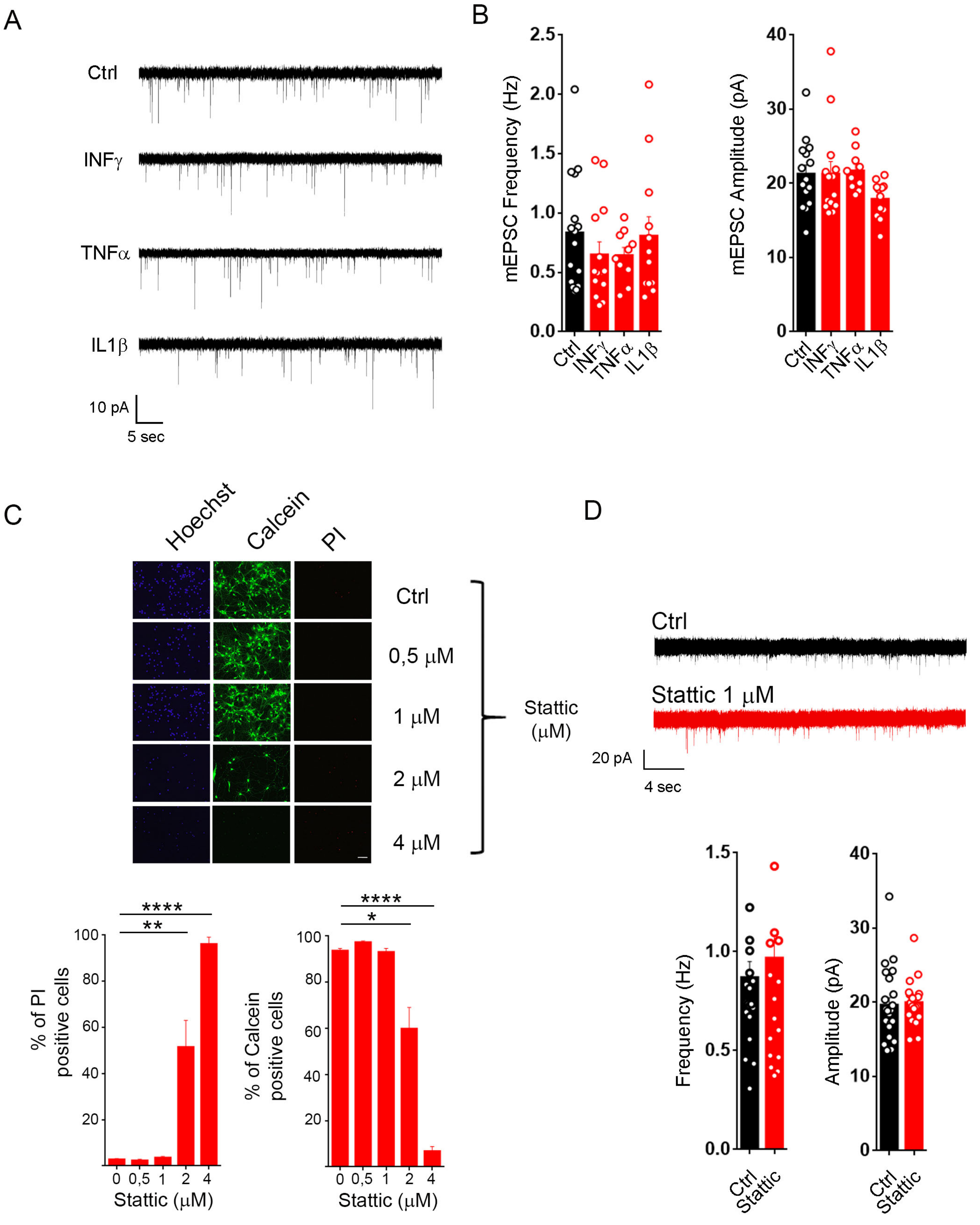


Mirabella et al., Figure S5

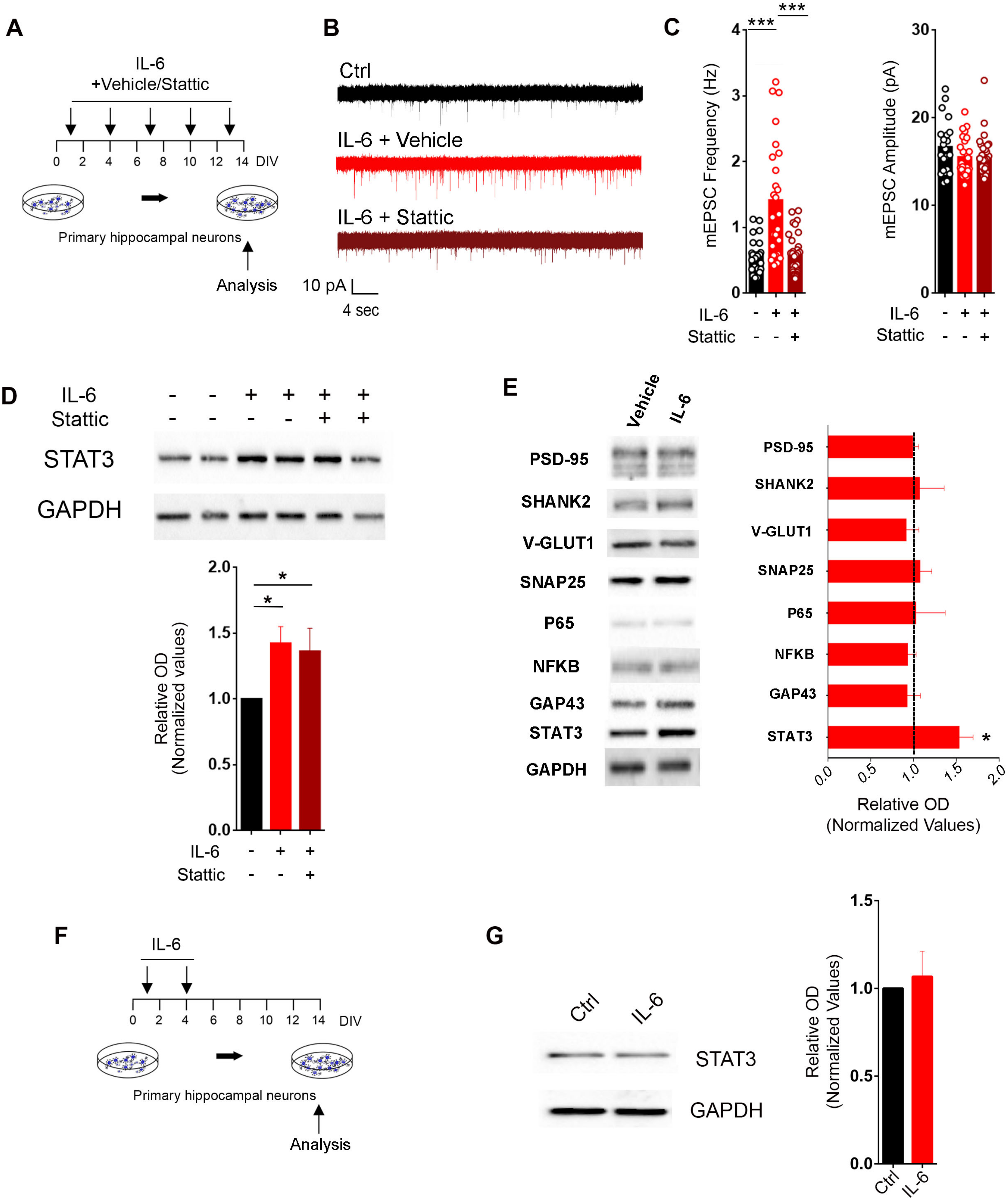


Mirabella et al., Figure S6

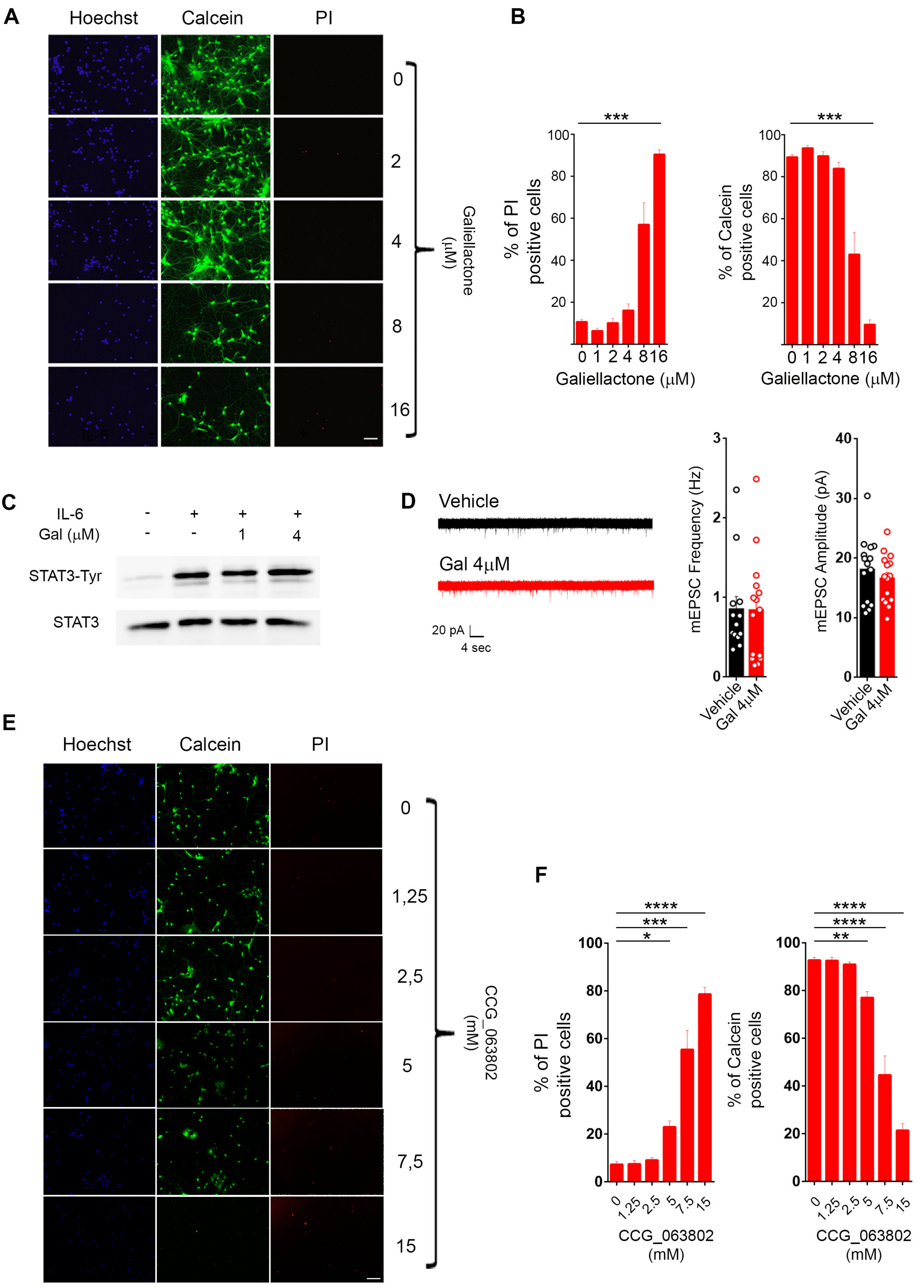


Mirabella et al., Figure S7

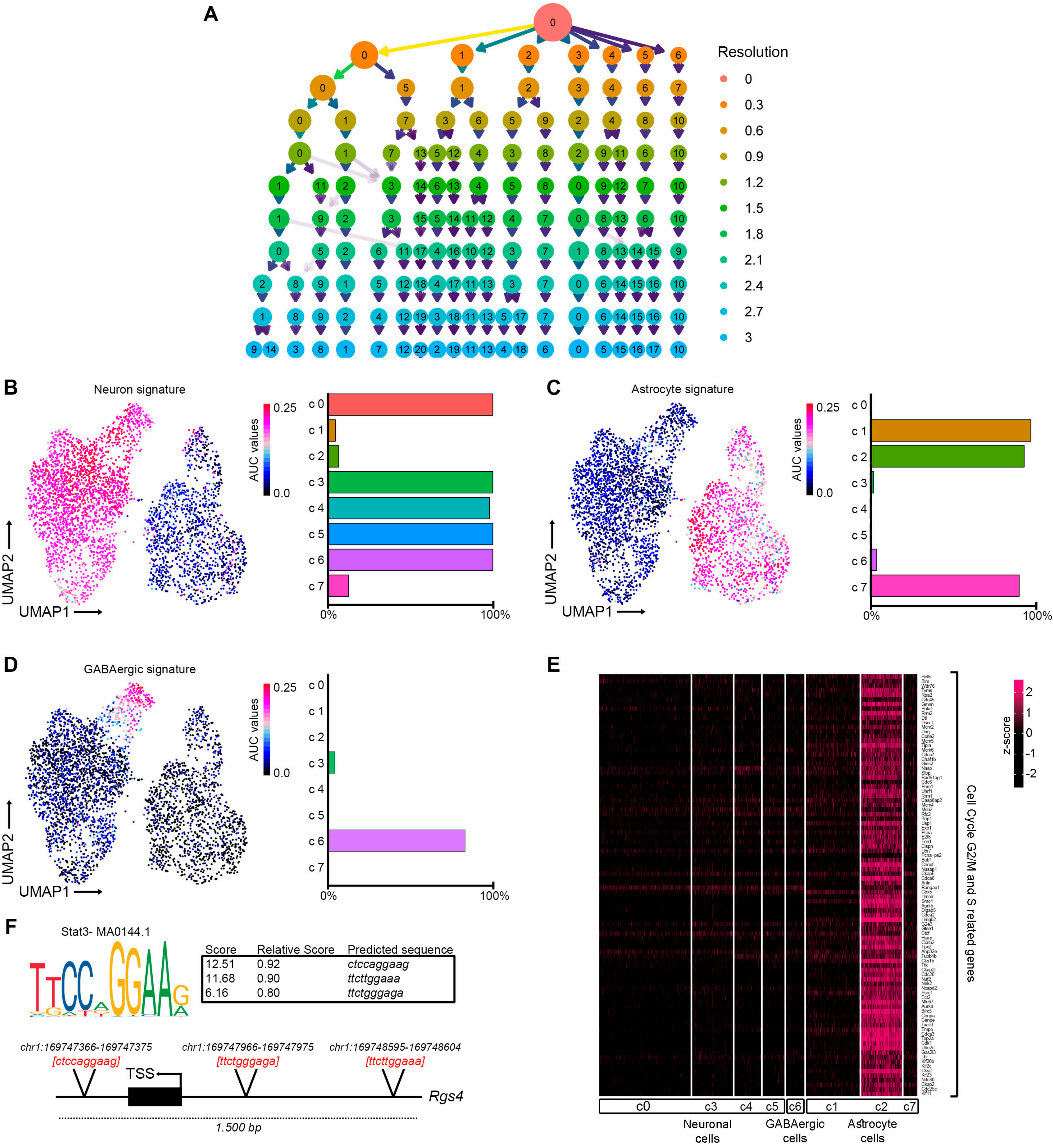


Mirabella et al., Figure S8
