## Supplementary material for "Transient Maternal IL-6 boosts glutamatergic synapses and disrupts hippocampal connectivity in the offspring": Table 1

**Table 1. List of gene signatures used for cluster identification**

| Neurons | Astrocyte | GABA | G2M | S |
|---|---|---|---|---|
| Snrpn | Edg1 | Gad2 | Bub1 | Blm |
| Gtl2 | Gja1 | Slc6a1 | Cenpf | Hells |
| LOC545465 | Gpr37l1 | Slc32a1 | Nusap1 | Wdr76 |
| Ryr2 | Gfap | Gad1 | Ckap5 | Tyms |
| Bex2 | Aqp4 | Htr3a | Cdca8 | Rpa2 |
| 3321401G04Rik | Atp1a2 | Dlx2 | Anln | Cdc45 |
| Calm2 | C1qa | Dlx6os1 | Rangap1 | Gmnn |
| Myo5a | C630015F21Rik | Dlx1 | Cbx5 | Pola1 |
| Clstn3 | Clu | Dlx5 | Hmmr | Rrm2 |
| Gria2 | Itgb5 | Lhx6 | Smc4 | Dtl |
| Camk2a | Mt1 | Rgs10 | Aurkb | Dscc1 |
| 2900011O08Rik | Pla2g7 | Tac1 | Dlgap5 | Mcm2 |
| 2900097C17Rik | Ppap2b | Sst | Cdca2 | Ung |
| Atp6v1g2 | Selpl | Npy | Hmgb2 | Ccne2 |
| Dpp6 | TC1568600 | Erbb4 | G2e3 | Mcm5 |
| Faim2 | Cst3 | Cck | Gtse1 | Tipin |
| Cyfip2 | Matn4 | Cxcl14 | Ctcf | Mcm6 |
| Gabarapl1 | Cd63 | Reln | Hjurp | Cdca7 |
| Gpr162 | Itgam | Ndnf | Ccnb2 | Chaf1b |
| Lba1 | Glul | Lypd1 | Tpx2 | Gins2 |
| Lhx9 | Cldn5 | Lmo1 | Anp32e | Nasp |
| Madd | Bcan | Chrm2 | Tubb4b | Slbp |
| Napb | Prdx6 | Sema5a | Cks1b | Rad51ap1 |
| Ndrg4 | Gldc | C1ql1 | Ttk | Cdc6 |
| Prkar1b | Mfge8 | Pvalb | Ckap2l | Prim1 |
| Ptprn | Magi1 | Pnoc | Cdc20 | Uhrf1 |
| Rtn1 | Mlph | Vip | Nuf2 | Rrm1 |
| Snap25 | Slc6a20 | Ntng1 | Nek2 | Casp8ap2 |
| Syp | Stab1 | Nos | Ncapd2 | Mcm4 |
| TC1430156 | Atp1a4 | | Psrc1 | Msh2 |
| Kifc2 | C030039L03Rik | | Ect2 | Rfc2 |
| Pacsin1 | Aldoc | | Mki67 | Brip1 |
| C030018G13Rik | Apoe | | Aurka | Usp1 |
| Reps2 | Dbi | | Birc5 | Exo1 |
| Prkaca | Lyzs | | Cenpa | Pcna |
| Map2k4 | Cspg3 | | Cenpe | E2f8 |
| D12Ertd553e | Dip2 | | Tacc3 | Fen1 |
| Alcam | Idb3 | | Tmpo | Clspn |

| | | | | |
|---|---|---|---|---|
| Disp2 | Sparc | | Cdca3 | Ubr7 |
| Map2k1 | 2900042E01Rik | | Top2a | Pcna-ps2 |
| Rab6 | Emx2 | | Cdk1 | |
| Add2 | Gsn | | Ube2c | |
| Chgb | Sept4 | | Gas2l3 | |
| Cplx1 | Sox8 | | Lbr | |
| Cx3cl1 | Sparcl1 | | Kif20b | |
| E130013N09Rik | Id3 | | Kif2c | |
| Gap43 | | | Cks2 | |
| Gria4 | | | Kif23 | |
| Mm.40569 | | | Ndc80 | |
| Ndfip1 | | | Ckap2 | |
| Nsg2 | | | Cdc25c | |
| Sh3gl2 | | | Kif11 | |
| Slc22a17 | | | | |
| Smarca2 | | | | |
| Syngr3 | | | | |
| TC1561993 | | | | |
| Uchl1 | | | | |
| Ywhag | | | | |
| Syt1 | | | | |
| Chn1 | | | | |
| Nefl | | | | |
| Camk2b | | | | |
| Grin2b | | | | |
| N28178 | | | | |
| Tuba4 | | | | |
| Gria3 | | | | |
| Jakmip1 | | | | |
| Gpr51 | | | | |
| Lrp11 | | | | |
| Chst1 | | | | |
| Egr1 | | | | |
| Fbxw5 | | | | |
| Flywch1 | | | | |
